# Generation of iPSC-derived CD4^+^ Th1 cells enhancing chimeric antigen receptor-T cell cytotoxicity

**DOI:** 10.1101/2025.04.24.650471

**Authors:** Yoshiki Furukawa, Midori Ishii, Shintaro Kinoshita, Ayaka Goto, Norihide Izumi, Kosuke Matsuzaki, Kensuke Yanashima, Tokuko Toyota, Kota Tachibana, Jun Ando, Hiromitsu Nakauchi, Miki Ando

## Abstract

CD4^+^ T cells are anticipated to enhance the overall immune response, including the anti-tumor activity of chimeric antigen receptor (CAR)-T cell therapy. In the past, we established a culture system to generate CD8^+^ T cells from iPS cells (iPSCs); however, it was challenging to generate CD4^+^ T cells. Drawing inspiration from the observation that adult T cell leukemia (ATL) cells are consistently CD4^+^ and possess Treg characteristics, we successfully generated CD4^+^ Treg cells by reprogramming ATL cells into iPSCs and then differentiating them into T cells. Gene expression analysis of this generation system suggested that *RUNX3* serves as a key regulator in T cell differentiation within the *ex vivo* generation system. By knocking out *RUNX3*, we demonstrated the generation of antigen-specific CD4^+^ Th1 cells via the iPSC route, thereby enhancing the activity of CD8^+^ CAR-T cells against GD2-expressing lymphoma. These technologies hold significant promise for contributing to “off-the-shelf” immunotherapies against malignant tumors, including solid tumors.

## Introduction

iPS cell (iPSC)-derived functionally rejuvenated antigen-specific cytotoxic T lymphocytes (rejuvenated CTLs; rejTs) generated from exhausted CTLs have shown promise in resolving the problem of T cell exhaustion ^1^. We have demonstrated that these rejTs have younger memory phenotypes, such as stem cell memory and central memory, and persist long term *in vivo*, resulting in continuous potent anti-tumor effect in refractory tumors ^2–4^. Moreover, we generated chimeric antigen receptor (CAR) combination rejTs (CARrejTs) that can target two antigens: one on the cell surface via the CAR and another intracellularly through the native T cell receptor (TCR). These CARrejTs exhibited strong antitumor effect *in vivo* against Epstein-Barr virus (EBV)-associated lymphomas ^5^ and against small cell lung cancer ^6^. Furthermore, seeking the capacity to administer rejTs rapidly to a large patient population, we edited HLA class-I antigens by CRISPR/Cas9 to suppress allogeneic immune rejection ^7^.

Interestingly, under our differentiation protocol iPSCs naturally differentiate into CD8 single positive (SP) cells mixed with double negative (DN) cells and not into CD4 SP cells. Even when iPSCs are established from a CD4^+^ T cell clone, differentiating them does not generate CD4 SP cells but instead CD8 SP cells ^1^. Many research groups have attempted to generate iPSC-derived CD4 SP cells. Ueda et al. generated CD4^+^ helper-like cells by forced expression of *CD4* retrovirally transduced into iPSC-derived T cells ^8^. However, if these were SP cells and if they indeed could enhance the cytotoxicity of effector T cells is unknown. Montel-Hagen et al. have reported organoid-induced differentiation of conventional T cells, among them a small contingent of CD4 SP cells ^9^ that may not yet be ready for therapeutic use. Yano et al. also generated CD4^+^ regulatory T cell-like cells using an artificial organoid system. The generated cells suppressed graft-versus-host disease *in vivo*. However, issues such as quality and stability of cell generation remained: A mixture of double negative, double positive, CD8 SP, and CD4 SP cells was produced, with positive selection required to extract CD4 SP cells ^10^. Stable generation of CD4 SP cells is still challenging.

CD4^+^ helper T cells have critical roles in various types of immunotherapy. For example, regulatory CD4^+^ T (Treg) cells have proven therapeutic efficacy in autoimmune disease such as systemic lupus erythematosus, inflammatory bowel disease, multiple sclerosis, and type 1 diabetes mellitus. Treg cells have also been proven to suppress allograft rejection in solid organ transplantation and graft-versus-host disease in hematopoietic stem-cell transplantation ^11, 12^. CD4^+^ Th1 cells have another important role in promoting the effector functions of CD8^+^ CTLs to develop and to sustain effective anti-tumor immunity ^13, 14^. CD4^+^ Th1 cells participate in the anti-tumor immune response by directly eliminating tumor cells or by providing cytokines and co-stimulatory signals that improve CD8^+^ CTL response efficacy ^13^. Moreover, synergistic efficacy against B-cell malignancies was demonstrated for CD4^+^ chimeric antigen receptor T (CAR-T) cells mixed with CD8^+^ CAR-T cells ^15^. CD4^+^ T cells generated from iPSCs would accordingly be useful for various types of immunotherapy, with stable generation of iPSC-derived CD4 SP cells an unlimited source of therapeutic CD4^+^ T cells for “off-the-shelf” therapy.

Our goal in this study was stable generation of iPSC-derived CD4 SP cells that could be used for clinical application. To generate CD4 SP cells, we first focused on adult T cell leukemia (ATL), because human T cell leukemia virus-1 (HTLV-1) infected CD4^+^ T cells clonally expand. We reprogrammed ATL cells into iPSCs, then differentiated these into T cells using our established method ^1, 16, 17^. Intriguingly, we succeeded in natural generation of CD4 SP cells from iPSCs.

We performed single-cell RNA sequencing (scRNA-seq) of iPSC-derived CD4 SP and CD8 SP cells to understand our success in CD4 SP cell differentiation and discovered several genes that might be key regulators of CD4/CD8 T cell lineage choice. We then used CRISPR/Cas9 gene editing technology to knock out (KO) these candidate genes and thereby to elucidate mechanisms of CD4/CD8 T cell lineage choice.

## Results

### CD4^+^ T cell clone-derived iPSCs differentiate into CD8 SP cells

First, we cloned CD4^+^ T cells from a healthy donor and reprogrammed the CD4^+^ T cell clone into iPSCs (CD4^+^ T cell clone-derived iPSCs; CD4-iPSCs). We next differentiated CD4-iPSCs originating from the CD4^+^ T cell clone into rejTs via our established method ^1, 16, 17^ (Figure 1A and 1B). Using flow cytometry, we analyzed CD4/CD8 expression on differentiated hematopoietic progenitor cells extracted from iPSC sacs (Week 2) to investigate what types of T cells CD4-iPSCs would emerge. The hematopoietic progenitor cells were initially CD4/CD8 double negative (DN) (Week 3) and then slightly trended towards CD4/CD8 double positive (DP) status (Week 5). They finally became CD8 SP after Week 6 and remained CD8 SP cells after T cell receptor (TCR) stimulation (CD8rejTs) (Figure 1C). We thus confirmed that even when generated from CD4 SP cells, iPSCs naturally differentiated into CD8 SP cells.

**Figure 1.**
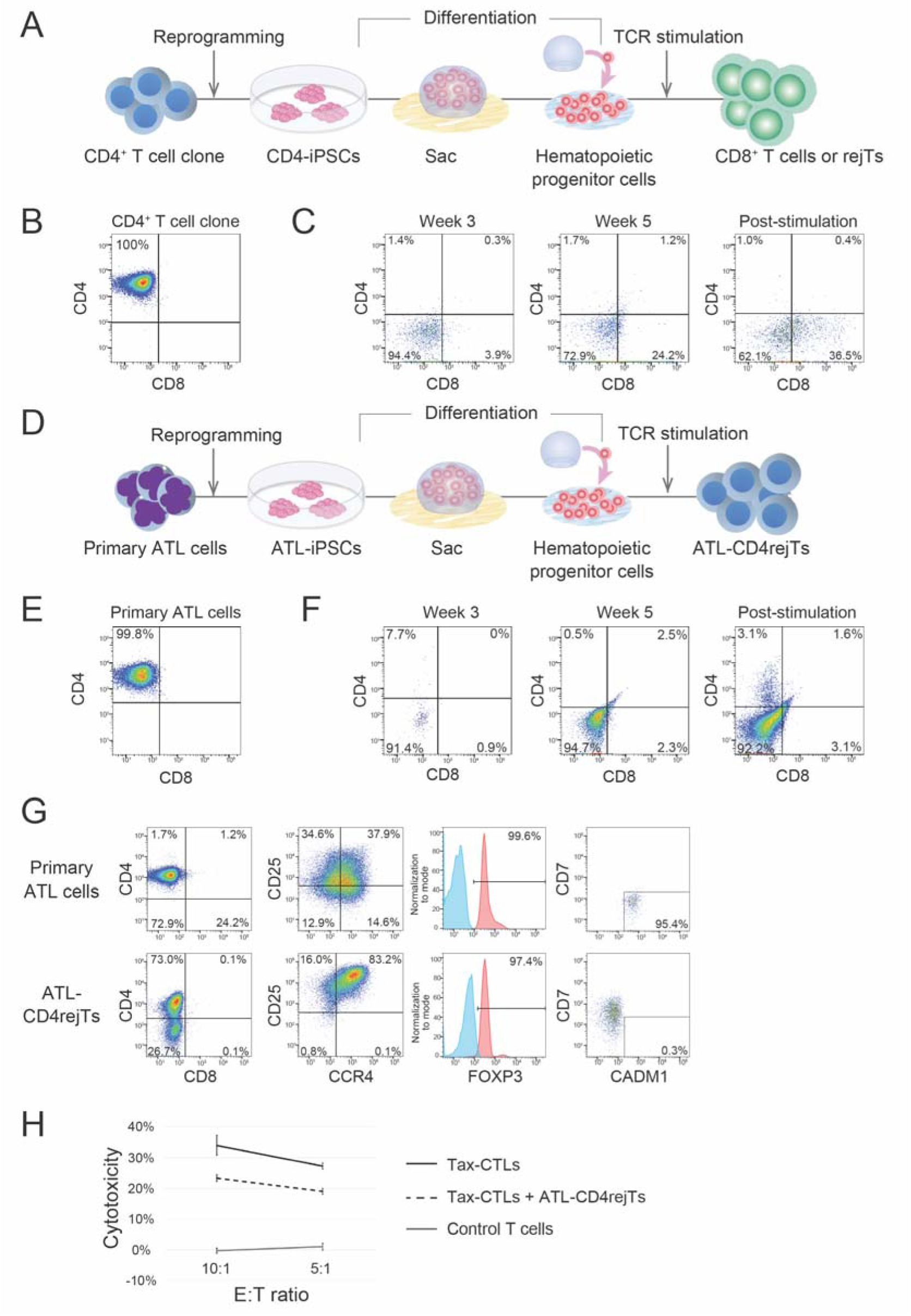
ATL-iPSCs differentiated into CD4 SP cells (A) Schematic illustration of differentiation of CD4-iPSCs. (B) Flow cytometric analysis of CD4^+^ T cell clone from a healthy donor. (C) Flow cytometric analysis of CD4-iPSCs differentiation from Week 3 when cells were transferred onto DL1/4-expressing C3H10T1/2 feeder cells. Cells were stimulated with PHA-L and irradiated PBMCs in CTL medium supplemented with 1% PSG in the presence of IL-2 PBMC (Week 6). Flow cytometry was performed every week until T cell receptor (TCR) stimulation (post-stimulation). The data shown are representative of at least three independent generations of CD4-iPSCs. (D) Schematic illustration of differentiation of ATL-iPSCs. (E) Flow cytometric analysis of primary ATL cells. (F) Flow cytometric analysis of ATL-iPSC differentiation from Week 3 when cells were transferred onto DL1/4-expressing C3H10T1/2 feeder cells. Cells were stimulated with PHA-L and irradiated PBMCs in NS-A2 CTL medium supplemented with 1% PSG in the presence of IL-2 (Week 6). Flow cytometry was performed every week until post-stimulation. The data shown are representative of at least three independent generations of ATL-iPSCs. (G) Characterization of primary ATL cells (top) and ATL-CD4rejTs (bottom) by flow cytometry. (H) *In vitro* ^51^Cr assay with Tax-CTL and Tax-CTL + ATL-CD4rejTs as effectors against primary ATL cells as targets. The E:T ratio was 10:1 and 5:1. Representative data of at least three independent experiments. Error bars represent ±SD.

### ATL-derived iPSCs differentiate into CD4 SP cells

In attempting to generate CD4 SP cells from iPSCs, we focused on ATL because HTLV-1 infected CD4^+^ T cells proliferate monoclonally in enormous numbers. We reprogrammed primary ATL cells (CD4 SP) from an ATL patient (Patient 1) to establish ATL-iPSCs. We differentiated these ATL-iPSCs using our established method ^1, 16, 17^ (Figure 1D and 1E). After extraction of hematopoietic progenitor cells from iPSC sacs (Week 2), we analyzed expression of CD3, CD4, and CD8 on these cells by flow cytometry to observe CD4/CD8 ratios during differentiation, with TCR stimulation terminally (Week 6) (Figure 1D). One week after hematopoietic stem cell extraction (Week 3), DN cells were found. These trended toward DP cells (Week 5), and subsequently yielded CD4 SP cells (ATL-CD4rejTs) mixed with DN cells. CD4 positive bead selection concluded generation of CD4 SP cells from iPSCs (Figure 1F).

We assessed reproducibility by generating several ATL-iPSC lines reprogrammed with primary ATL cells from the same patient (Patient 1) as well as from a different patient (Patient 2) with acute ATL (Supplementary Figure 1A). All ATL-iPSCs efficiently differentiated into CD4 SP cells and robustly proliferated.

### ATL-CD4rejTs showed a regulatory T cell phenotype

To characterize ATL-CD4rejTs we analyzed them by flow cytometry. We compared the phenotype of these T cells with that of primary ATL cells. Substantially most ATL-CD4rejTs were CD4^+^, CD25^+^, CCR4^+^, and FOXP3^+^, of Treg phenotype, the same as that of the primary ATL cells (Figure 1G). The cells differed in CADM1 expression, used to define ATL cells ^18–20^. Primary ATL cells expressed CADM1 and ATL-CD4rejTs did not, evidence that ATL-CD4rejTs were not ATL cells (Figure 1G).

To investigate if Treg-phenotype ATL-CD4rejTs exert a suppressive effect on effector T cells, we conducted chromium-51 (^51^Cr) release assays, measuring the cytotoxicity of HTLV-1 Tax-specific CTLs (Tax-CTLs), with or without ATL-CD4rejTs, against primary ATL cells. At an effector : target (E:T) ratio of 10:1, 34 ± 3.2% of ATL cells were lysed by Tax-CTLs. On adding ATL-CD4rejTs, cytotoxicity fell to 23.4 ± 1.0%, demonstrating ATL-CD4rejTs to exert a suppressive effect (Figure 1H).

We then, using qPCR, measured expression levels of HTLV-1 oncogenic factors *Tax* and *HBZ* in primary ATL cells, ATL-iPSCs, and ATL-CD4rejTs. Expression of *Tax* relative to primary ATL cells was 0.00003 : 1 in ATL-iPSCs and 0.0072 : 1 in ATL-CD4rejTs. Likewise, relative expression of *HBZ* relative to primary ATL cells was 0.0004 : 1 in ATL-iPSCs and 0.0710 : 1 in ATL-CD4rejTs (Supplementary Figure 1B). ATL-CD4rejTs thus expressed *Tax* and *HBZ* at much lower levels than did primary ATL cells.

To investigate generated ATL-CD4rejTs further, we performed scRNA-seq and cellular indexing of transcriptomes and epitopes (CITE) sequencing (CITE-seq) of both primary ATL cells and ATL-CD4rejTs (Supplementary Figures 1C and 1D). RNA and surface antigen cocktail antibody barcoding-based clustering of the CITE-seq dataset yielded 9 individual clusters. Volcano plotting revealed that the most significant differentially-expressed gene (DEG) in primary ATL cells was *GNLY*, while the most significant DEG in ATL-CD4rejTs was *KLRB1*. Thus a clear difference was demonstrated between primary ATL cells and ATL-CD4rejTs regarding which genes were highly expressed (Supplementary Figure 1E and 1F).

### *RUNX3* was highly expressed in CD8rejTs but not in ATL-CD4rejTs

To understand further the mechanism of CD4/CD8 lineage choice, we also compared gene expression in ATL-CD4rejTs with that in CD8rejTs, using scRNA-seq and CITE-seq. CD8rejTs generated from CD4-iPSCs (healthy donor CD4^+^ T cell clone-derived iPSCs) (Figure 1A and 1C) were used as a control. We performed uniform manifold approximation and projection (s) analysis and compared expression levels of genes associated with CD4/CD8 T cell lineage choice in CD8rejTs with those in ATL-CD4rejTs using violin plot analysis ^21–29^ (Figure 2A and Supplementary Figure 2A and 2B). The key transcription factor, known as Thpok or cKrox, that directs commitment to CD4 T cell lineage is encoded by *ZBTB7B* ^21–23^, while the key transcription factors that direct commitment to CD8 T cell lineage are encoded by *RUNX1* and *RUNX3* ^24–26^. In a murine model, *Socs1*, *Socs3*, and *Gimap5* also are reportedly related to CD4 T cell lineage choice ^28, 29^. In respect of these key transcription factor genes, we discovered that *RUNX3* was expressed in CD8rejTs at high levels, but in ATL-CD4rejTs at low levels. Although expression of *RUNX1*, *SOCS1*, *SOCS3*, and *GIMAP5* also differed substantially, *RUNX3* expression differed most prominently between CD8rejTs and ATL-CD4rejTs. Interestingly, *ZBTB7B*, reportedly the key regulatory gene that directs commitment to CD4 T cell lineage ^21–23^, was not expressed in either CD8rejTs or ATL-CD4rejTs (Figure 2A and Supplementary Figure 2B). Among known key regulatory genes affecting CD4/CD8 T cell lineage choice, then, *RUNX3* expression levels differed most between CD8rejTs and ATL-CD4rejTs.

**Figure 2.**
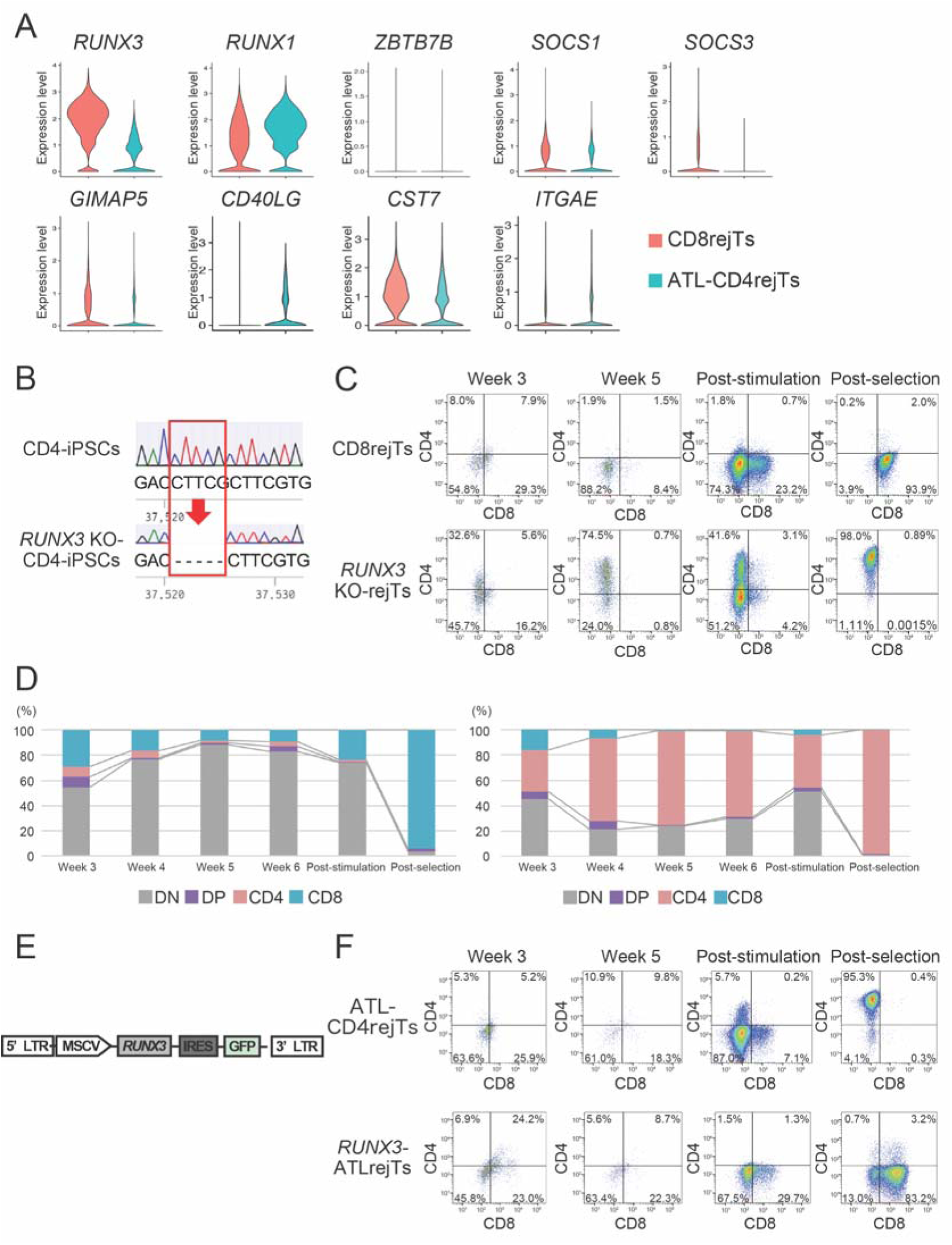
*RUNX3* was confirmed as key regulator in CD4/CD8 T cell lineage choice (A) Violin plots (right) of genes associated with CD4/CD8 lineage choice (*RUNX3*, *RUNX1*, *ZBTB7B*, *SOCS1*, *SOCS3*, *GIMAP5, CD40LG, CST7, ITGAE*). (B) *RUNX3* KO in healthy donor CD4^+^ T cell clone derived-iPSCs using CRISPR/Cas9 gene editing technology. Five base pairs were deleted, causing a frame shift mutation with gain of a stop codon in exon 5. (C) Flow cytometric analysis of CD8rejTs and *RUNX3* KO-rejTs differentiation from iPSC-derived hematopoietic progenitor cells (Week 3) when cells were transferred onto DL1/4-expressing C3H10T1/2 feeder cells. Cells were stimulated with PHA-L and irradiated PBMCs (Week 6). Flow cytometry was performed every week until after T cell receptor (TCR) stimulation (post-stimulation). The data shown are representative of at least three independent generations of CD8rejTs and *RUNX3* KO-rejTs. (D) Weekly comparison of the percentage of CD4 SP cells and CD8 SP cells within CD8rejTs and *RUNX3*-KO-rejTs from Week 3 of differentiation until post-stimulation. The data shown are representative of at least three independent generations of CD8rejTs and *RUNX3* KO-rejTs. (E) Schema, lentiviral vector construct used for overexpression of *RUNX3*. (F) Flow cytometric analysis of ATL-CD4rejTs and *RUNX3*-overexpressing ATLrejTs (*RUNX3-*ATLrejTs) differentiation (Week 3) when cells were transferred onto DL1/4-expressing C3H10T1/2 feeder cells. Cells were stimulated with PHA-L and irradiated PBMC (Week 6). Flow cytometry was performed every week until post-stimulation. The data shown are representative of at least three independent generations of ATL-CD4rejTs and *RUNX3*-ATLrejTs.

### *RUNX3* KO iPSCs differentiated into CD4 SP cells

As *RUNX3* expression was markedly higher in CD8rejTs differentiated from CD4-iPSCs than in ATL-CD4rejTs (Figure 2A), we hypothesized that on knocking out *RUNX3* in CD4-iPSCs these would differentiate into CD4 SP cells instead of CD8 SP cells. Therefore, we knocked out *RUNX3* in CD4-iPSCs using CRISPR/*Cas9* gene editing technology. A deletion of 5 base pairs resulted in a frameshift mutation with gain of a stop codon in exon 5 and yielded *RUNX3* KO cells (Figure 2B).

We differentiated these *RUNX3*-KO-CD4-iPSCs using our established method ^1, 16, 17^. We also differentiated wild-type (WT) CD4-iPSCs as a control for CD4/CD8 percentages during differentiation. After extraction of hematopoietic progenitor cells from *RUNX3*-KO-CD4-iPSC sacs (Week 2), we weekly examined expression of CD3, CD4, and CD8 on *RUNX3*-KO-rejTs by flow cytometry. The percentage of CD4 SP tended to increase over time in *RUNX3*-KO-rejTs, while the percentage of CD8 SP cells tended to decrease. In WT CD4-iPSCs, the percentage of CD4 SP tended to decrease over time, while the percentage of CD8 SP cells tended to increase (Figure 2C and 2D). Thus with *RUNX3* KO, CD4-iPSCs successfully differentiated into CD4 SP cells instead of CD8 SP cells.

### *RUNX3*-overexpressing ATL-iPSCs differentiated into CD8 SP cells

To learn if iPSCs would generate CD8 SP cells with overexpression of *RUNX3*, we overexpressed *RUNX3* in ATL-iPSCs (*RUNX3*-ATL-iPSCs) that without manipulation differentiated into CD4 SP cells. After we transduced retroviral *RUNX3* (Figure 3E) into ATL-iPSCs, we differentiated ATL-iPSCs and *RUNX3*-ATL-iPSCs at the same time using our established method ^1, 16, 17^ to generate ATL-CD4rejTs and *RUNX3*-ATL-rejTs respectively and compared CD4/CD8 percentages during differentiation (Figure 3F). After Week 2, we weekly performed flow cytometry to investigate CD3, CD4, and CD8 expression on differentiated cells every week. In ATL-iPSCs the percentage of CD8 SP tended to decrease over time and the percentage of CD4 SP cells tended to increase. The percentage of CD8 SP tended to increase over time in *RUNX3*-ATL-rejTs and the percentage of CD4 SP cells decreased. These findings demonstrated that with overexpression of *RUNX3* ATL-iPSCs differentiated into CD8 SP cells instead of CD4 SP cells and established that *RUNX3* is a key regulator of CD4/CD8 T cell lineage choice in our iPSC-derived T cell differentiation system.

**Figure 3.**
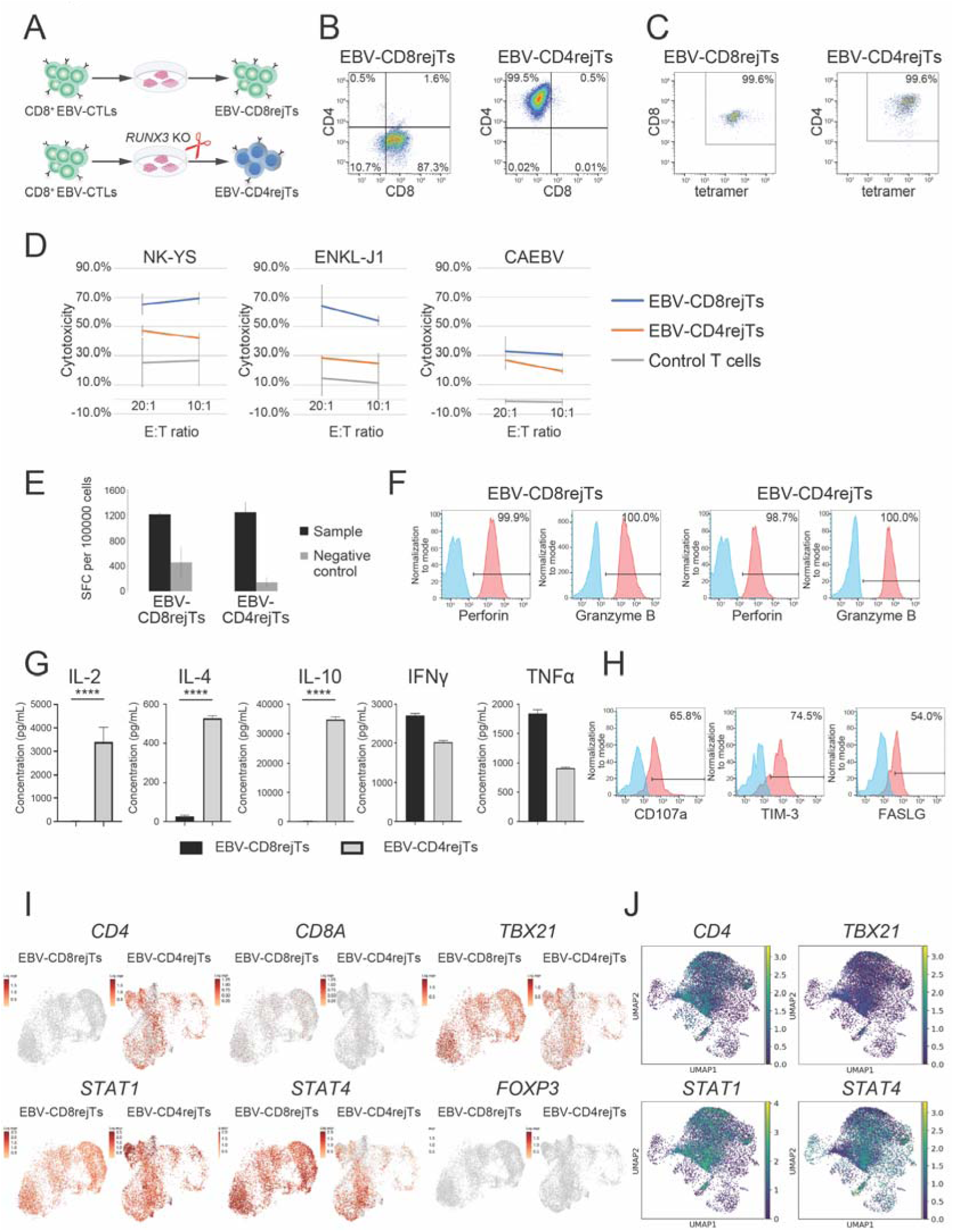
EBV-CD4rejTs maintained MHC class I restriction and exhibited antigen-specific cytotoxic activity (A) Schematic illustration of generation of EBV-CD8rejTs (top) and EBV-CD4rejTs (bottom). (B) Flow cytometric CD4/CD8 analysis of post-selected EBV-CD8rejTs and EBV-CD4rejTs. (C) Flow cytometric EBV LMP2-tetramer analysis of post-selected EBV-CD8rejTs and EBV-CD4rejTs. (D) *In vitro* ^51^Cr assay with EBV-CD8rejTs and EBV-CD4rejTs as effectors against NK-YS cells (left), ENKL-J1 cells (middle), and primary CAEBV cells (right) as targets. The effector-to-target (E:T) ratios were 20:1 and 10:1. Representative data of at least three independent experiments. Error bars represent ±SD. (E) IFN-γ Elispot in response to stimulation with peptide pulse. Negative control shows IFN-γ release in response to no peptide pulse. Results are expressed as spot-forming cells (SFC) per 1.0 x 10^5^ cells. Representative data of at least three independent experiments. Error bars represent ±SD. (F) Histograms of fluorescence intensity of expression of intracellular staining of perforin and granzyme B for EBV-CD8rejTs and EBV-CD4rejTs (red). Negative controls (fluorescence-minus-one control) are shown in blue. (G) Cytometric bead array for measuring cytokines (IFNγ, TNFα, IL-4, IL-10 and high-sensitivity IL-2) produced by EBV-CD8rejTs and EBV-CD4rejTs after 24 of coculture with NK-YS cells. Error bars represent ± SEM. ****p < 0.0001, one-way ANOVA. (H) Histograms of fluorescence intensity of expression of surface markers related to Th1 cells (CD107a, CD366, FASLG) for EBV-CD8rejTs and EBV-CD4rejTs (red). Negative controls (fluorescence-minus-one controls) are shown in blue. (I) UMAPs of genes associated with Th1 cells (*CD4*, *CD8A*, *TBX21*, *STAT1*, *STAT4*). (J) Overlapping of EBV-CD4rejTs scRNA-seq dataset with a conventional T cell scRNA-seq dataset. EBV-CD4rejTs scRNA-seq data are shown in orange and conventional T cells data are shown in blue. UMAPs of overlapping of *CD4*, *TBX21*, *STAT1*, and *STAT4* are presented.

### EBV-CD4rejTs maintained antigen-specific cytotoxicity after *RUNX3* KO

We extended these investigations by knocking out *RUNX3* in iPSCs originated from antigen-specific CTLs to examine if *RUNX3* KO-iPSC-derived CTLs also differentiated into CD4 SP cells. We knocked out *RUNX3* in iPSCs derived from EBV latent membrane protein (LMP2)-specific CD8^+^ CTLs (EBV-CTLs) (Figure 3A). Although EBV-CTL-derived iPSCs (EBV-iPSCs) always differentiate into CD8^+^ T cells (EBV-CD8rejTs) ^2^, we hypothesized that with *RUNX3* KO in EBV-iPSCs, CD4 SP cells that maintain LMP2 antigen specificity could be generated. Interestingly, these generated EBV-rejTs were CD4 SP cells (EBV-CD4rejTs) (Figure 3B) but maintained major histocompatibility complex (MHC) class I-restricted LMP2 antigen-specificity (Figure 3C). To investigate the function of EBV-CD4rejTs, we conducted ^51^Cr release assays. The cytotoxicity of EBV-CD8rejTs and EBV-CD4rejTs against EBV-associated lymphoma cells was measured. At an E:T ratio of 20:1, 61.3 ± 2.9% of NK-YS cells were lysed by EBV-CD8rejTs and 19.0 ± 2.5% of NK-YS cells were lysed by EBV-CD4rejTs (Figure 3D, left). Similarly, at an E:T ratio of 20:1, the percentages of ENKL-J1 cells lysed by EBV-CD8 rejTs and EBV-CD4 rejTs were 64.3 ± 14.1% and 28.6 ± 1.1%, respectively (Figure 3D, middle). The percentages of primary chronic active EBV disease (CAEBV) cells lysed by EBV-CD8rejTs and EBV-CD4rejTs were 32.5 ± 9.9% and 26.8 ± 6.5%, respectively (Figure 3D, right). Therefore, although weaker than EBV-CD8rejTs, EBV-CD4rejTs exhibited class I-restricted LMP2 antigen-specific cytotoxicity.

We measured IFN-γ levels for EBV-CD8rejTs and EBV-CD4rejTs with IFN-γ ELISPOT assays after stimulation with LMP2 peptide. Both EBV-CD8rejTs and EBV-CD4rejTs showed specific activity against LMP2 (1209.0 ± 24.04 and 1242.67 ± 73.08 IFN-γ spot-forming cells [SFCs]/10,000, respectively) (Figure 3E). We also evaluated the expression of intracellular perforin and granzyme B by flow cytometry. EBV-CD4rejTs expressed both perforin and granzyme B strongly (98.7% and 100.0%, respectively), similar to EBV-CD8rejTs (99.9% and 100.0%, respectively) (Figure 3F).

### EBV-CD4rejTs secreted numerous cytokines and showed Th1 phenotype

In further cytotoxic-effect mechanistic studies of EBV-CD4rejTs we used cytometric bead arrays to measure cytokine release levels. EBV-CD4rejTs secreted IL-2, IL-4, and IL-10 at significantly higher levels than did EBV-CD8rejTs (all, p<0.0001). Although at levels lower than EBV-CD8rejTs, EBV-CD4rejTs also secreted IFN-γ and TNF-α (Figure 3G). Using flow cytometry, we determined that EBV-CD4rejTs expressed the cell-surface markers CD107a, CD366, and FASLG (Figure 3H). We also conducted scRNA-seq of EBV-CD8rejTs and EBV-CD4rejTs and compared gene expressions. EBV-CD4rejTs significantly expressed *CD4* and both EBV-CD8rejTs and EBV-CD4rejTs clearly expressed *TBX21*, *STAT1*, and *STAT4* (Figure 3I).

To determine how similar EBV-CD4rejTs are to conventional CD4^+^ T cells, we compared scRNA-seq data for EBV-CD4rejTs with an existing dataset for peripheral blood T cells (Figure 3J and Supplementary Figure 3A-C) ^30^. Not only did EBV-CD4rejTs display a Th1 phenotype ^31^; they also overlapped with cells annotated as Th1 cells (*TBX21*, *STAT1*, *STAT4* positive cells) in the conventional-cell dataset (Figure 3J, Supplementary Figure 3B and 3C) ^32^. EBV-CD4rejTs thus proved to secrete numerous cytokines, including IL-2, IFN-γ, and TNF-α, and to express genes (*TBX21*, *STAT1*, and *STAT4*) and surface antigens (CD107a, CD366, and FASLG) in patterns consistent with the Th1 phenotype in conventional cells. We also succeeded in generating Th1 cells from by *RUNX3* KO human papilloma virus (HPV)-specific CTL-derived iPSCs (Supplementary Figure 3D - 3G).

### Knocking out *RUNX3* downregulated genes associated with CD8 T cell lineage choice

We performed RNA-seq in cells differentiated from CD8^+^ EBV-CTL-derived WT-iPSCs and KO-iPSCs into EBV-CD8rejTs and EBV-CD4rejTs, respectively, on days 0, 7, 14, 21, and 28 (Supplementary Figure 4). EBV-CD8rejTs did not express CD4 and EBV-CD4rejTs expressed it on day 28. By day 28 EBV-CD8rejTs expressed CD8A and CD8B while EBV-CD4rejTs did not. Knockout of *RUNX3* was confirmed in EBV-CD4rejTs. Among genes associated with CD4 T cell lineage choice, expression of *ZBTB7B* increased similarly in EBV-CD8rejTs and EBV-CD4rejTs. *CD40LG*, also associated with CD4 T cell lineage choice (27), was somewhat more highly expressed in EBV-CD8rejTs than in EBV-CD4rejTs, suggesting that expression of *ZBTB7B* and *CD40LG* did not directly affect CD4 differentiation. Among genes associated with CD8 T cell lineage choice, expression of *RUNX1, SOCS1*, and *SOCS3* was similar in EBV-CD8rejTs and EBV-CD4rejTs. Expression of *GIMAP5*, *ITGAE*, and *CST7* increased in EBV-CD8rejTs and was downregulated in *RUNX3* KO EBV-CD4rejTs ^27^ (Supplementary Figure 4).

With KO of *RUNX3* alone, genes associated with CD8 T cell lineage choice such as *GIMAP5*, *ITGAE*, and *CST7* were downregulated. These data suggest that *RUNX3* may be upstream of these 3 genes, strongly influencing CD4 differentiation in CD4/CD8 T cell lineage choice.

### EBV-CD4rejTs exhibited helper function with CD8^+^ CART cells and decreased the exhaustion level

To investigate whether Th1-phenotype EBV-CD4rejTs show helper functions, we cocultured EBV-CD4rejTs with disialoganglioside (GD2)-chimeric antigen receptor (CAR) T cells (GD2-CARTs). GD2-CARTs reportedly become exhausted due to tonic signaling ^6^ and are an ideal model in examining if EBV-CD4rejTs can improve effector function. CD8^+^/CAR^+^ sorted GD2-CARTs were cocultured with EBV-CD4rejT at a 1:1 ratio (mixed cells), without cytokines, to examine proliferative capacity and cytotoxicity. Compared with GD2-CARTs alone and EBV-CD4rejTs alone, mixed cells proliferated significantly more strongly, indicating that EBV-CD4rejTs (Th1 cells) assisted GD2-CARTs (GD2-CARTs *vs*. mixed cells; p= 0.0004) (Figure 4A). To examine the cytotoxicity of GD2-CARTs, EBV-CD4rejTs, and mixed cells against GD2^+^ EBV-associated lymphoma cells, we conducted ^51^Cr release assays. Cells were cultured without cytokines for 3 days before assay. The same number of cells was used for each experiment. On the day of assay, we evaluated CD4, CD8, and tetramer positivity in GD2-CARTs, EBV-CD4rejTs, and mixed cells by flow cytometry. We confirmed that GD2-CARTs were CD8 SP cells not having LMP2 antigen specificity, EBV-CD4rejTs were CD4 SP cells with MHC class I-restricted LMP2 antigen specificity, and mixed cells had both CD8 SP cells and CD4 SP cells with antigen specificity (Figure 4B). We also examined the cytotoxicity of GD2-CARTs, EBV-CD4rejTs, and mixed cells when cocultured with GD2^+^ ENKL-J1 cells at E:T ratios of 40:1, 20:1, 10:1, and 5:1. At an E:T ratio of 40:1, 25.6 ± 2.9% of ENKL-J1 cells were lysed by mixed cells, 14.7 ± 0.2% of ENKL-J1 cells were lysed by GD2-CARTs, and 14.1 ± 1.7% of ENKL-J1 cells were lysed by EBV-CD4rejTs (Figure 4C). By coculturing GD2-CARTs with EBV-CD4rejTs, a synergistic effect on cytotoxicity was achieved at E:T ratios of 40:1, 20:1, and 10:1 (all, p < 0.001) (Figure 4D).

**Figure 4.**
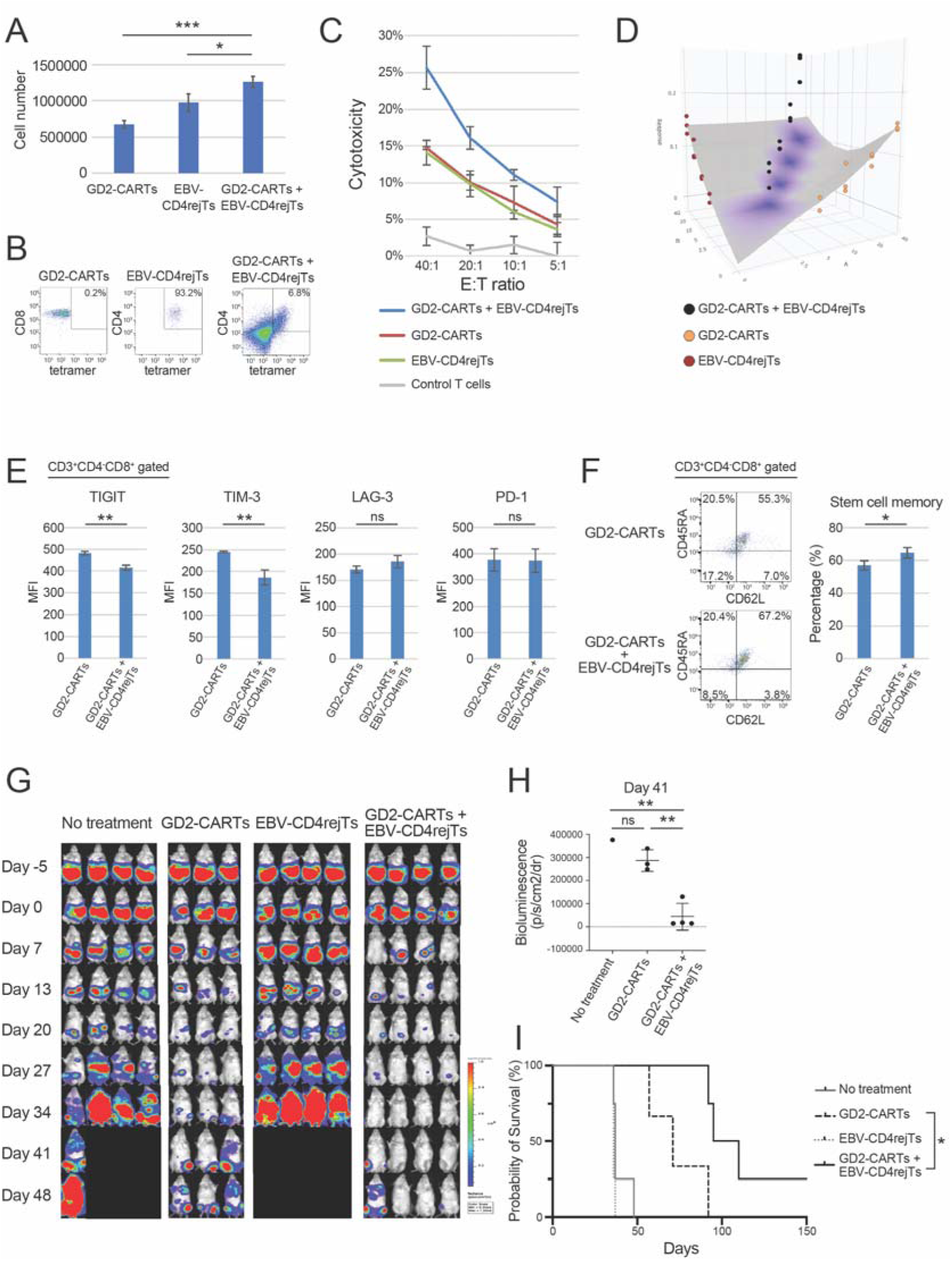
EBV-CD4rejTs exhibited helper function toward CD8^+^ CART cells *in vitro* and *in vivo* (A) Proliferation of cells used for *in vitro* ^51^Cr assay (CD8^+^/CART^+^ GD2-CARTs, EBV-CD4rejTs, and GD2-CARTs mixed with EBV-CD4rejTs) 2 days after 1.0 M cells were collected. Data represent means of at least 2 experiments (n=3). Error bars represent ± SD. *p < 0.05, ***p < 0.001, one-way ANOVA. (B) Flow cytometric tetramer analysis of cells used for *in vitro* ^51^Cr assay (GD2-CARTs, EBV-CD4rejTs, mixed cells). (C) *In vitro* ^51^Cr assay with GD2-CARTs, EBV-CD4rejTs, mixed cells, and control T cells as effectors against ENKL-J1 cells as targets. The E:T ratios were 40:1, 20:1, 10:1, and 5:1. Representative data of at least three independent experiments. Error bars represent ±SD. (D) By coculturing GD2-CARTs with EBV-CD4rejTs, synergistic effect was achieved at E:T ratios of 40:1, 20:1, and 10:1 (E) MFI of exhaustion markers (TIGIT, TIM-3, LAG-3, PD-1) of GD2-CARTs was compared with that of GD2-CARTs cocultured with EBV-CD4rejTs. Flow cytometry was performed 7 days after coculture with ENKL-J1 cells. Representative data of at least two independent experiments. Error bars represent ±SD. **p < 0.01, one-way ANOVA; ns, not significant. (F) Representative flow cytometrogram of memory phenotype among GD2-CARTs and GD2-CARTs cocultured with EBV-CD4rejTs. Flow cytometry was performed 7 days after coculture with ENKL-J1 cells. Representative data of at least two independent experiments. Error bars represent ±SD. *p < 0.05, unpaired Student’s *t* test (two-tailed). (F) *In vivo* bioluminescence imaging of mice engrafted with ENKL-J1 cells in untreated (control group, n = 4) and treated animals (GD2-CARTs group, n=3; EBV-CD4 rejTs group, n = 4, and GD2-CARTs mixed with EBV-CD4rejTs group n=4). Data represent at least 2 independent experiments. (G) Quantitation of tumor burden on day 41 is shown: No treatment group (*n* = 1), GD2-CARTs group (*n* = 3), and GD2-CARTs mixed with EBV-CD4rejTs (*n* = 4). *, p<0.05; **, p<0.01 by one-way ANOVA. (H) Kaplan–Meier curves representing percentage survival of groups: no treatment group (*n* = 4), GD2-CARTs group (*n* = 3), EBV-CD4rejTs group (n=3) and GD2-CARTs mixed with EBV-CD4rejTs (*n* = 4). *, p < 0.05 by log-rank test.

To evaluate the extent of exhaustion in EBV-CD4rejTs, we cocultured GD2-CARTs, EBV-CD4rejTs, and mixed cells with ENKL-J1 cells for 7 days. We examined exhaustion markers (TIGIT, TIM-3, LAG-3, and PD-1) on cocultured cells by flow cytometry on days 0 and 7 of coculture to see if coculture with EBV-CD4rejT affected the extent of GD2-CART cell exhaustion. The mean fluorescence intensity (MFI) of TIGIT in GD2-CARTs was significantly higher than that in mixed cells (482.3 ± 8.4 *vs*. 414.7 ± 12.0, p=0.0013). The MFI of TIM-3 in GD2-CARTs was also significantly higher than that in mixed cells (245.0 ± 2.0 *vs*. 187.0 ± 17.6, p=0.0045). LAG-3 and PD-1, however, did not differ significantly between GD2-CARTs and mixed cells (p=0.1238 and p=0.9376, respectively) (Figure 4E).

We also analyzed memory phenotype subsets based on expression of the cell-surface markers CD45RA and CD62L. The percentage of stem cell memory T cells (CD45RA^+^, CD62L^+^) among GD2-CARTs significantly increased when cocultured with EBV-CD4rejTs (57.1±2.86% *vs*. 63.8 ± 3.21%, p=0.0357) (Figure 4F).

Coculture of GD2-CARTs with EBV-CD4rejTs improved the proliferation and cytotoxicity of GD2-CARTs. Furthermore, expression of exhaustion markers decreased and proportional stem cell memory increased in CD8^+^ GD2-CARTs. These results demonstrated that Th1-phenotype EBV-CD4rejTs support effector function of GD2-CART cells *in vitro*.

### EBV-CD4rejTs enhanced the cytotoxicity of GD2-CARTs *in vivo*

We used the cocultured cells from *in vitro* experiments to evaluate whether EBV-CD4rejTs also exhibited helper function *in vivo*. We intraperitoneally injected FFluc-labelled ENKL-J1 cells (1 × 10^6^ cells) into NOG mice. The cells’ bioluminescence was monitored as an indicator of tumor growth. Five days after tumor inoculation, mice were divided into an untreated group (control) and 3 treatment groups in which mice were intraperitoneally treated with GD2-CARTs, EBV-CD4rejTs, or mixed cells (4.0 x 10^6^ cells each) (Figure 4F). GD2-CARTs and EBV-CD4rejTs were cocultured at a 1:1 ratio for 3 days before injection. Bioluminescence gradually increased in untreated mice and in EBV-CD4rejT treated mice. In contrast, tumor signal was suppressed in both GD2-CART treated mice and mixed-cell treated mice (Figure 4G). The tumor signal in mice treated with mixed cells was significantly lower than in mice treated with GD2-CARTs on day 41 (286000 ± 47318.07 *vs*. 44025 ± 57333.026, p=0.0046; Figure 4H). Moreover, mice treated with mixed cells survived significantly longer (92 - 210 days) than mice treated with GD2-CARTs (57 - 92 days, p= 0.0288) (Figure 4I).

Coculture with EBV-CD4rejTs conferred on GD2-CARTs greater tumor suppression and prolonged mouse survival. EBV-CD4rejTs supported GD2-CARTs Th1 helper function not only *in vitro*, but also *in vivo*.

### Successful generation of CD4^+^ CTLs from MHC class II-restricted CD4**^+^**CTL-derived iPSCs

CD4^+^ CTLs represent a rare subpopulation of MHC class II-restricted CD4^+^ T cells with cytotoxic activity, achieved through the release of perforin and granzyme B. They may also exhibit characteristics such as the ability to produce IFNγ, either alone or in combination with other cytokines like TNFα and IL-2, which overlap with the features of Th1 cells ^33, 34^. Thus, we investigated whether we could generate MHC class II-restricted rejuvenated CD4^+^ CTLs from *RUNX3* KO iPSCs established from CD4^+^antigen-specific CTLs (Figure 5A).

**Figure 5.**
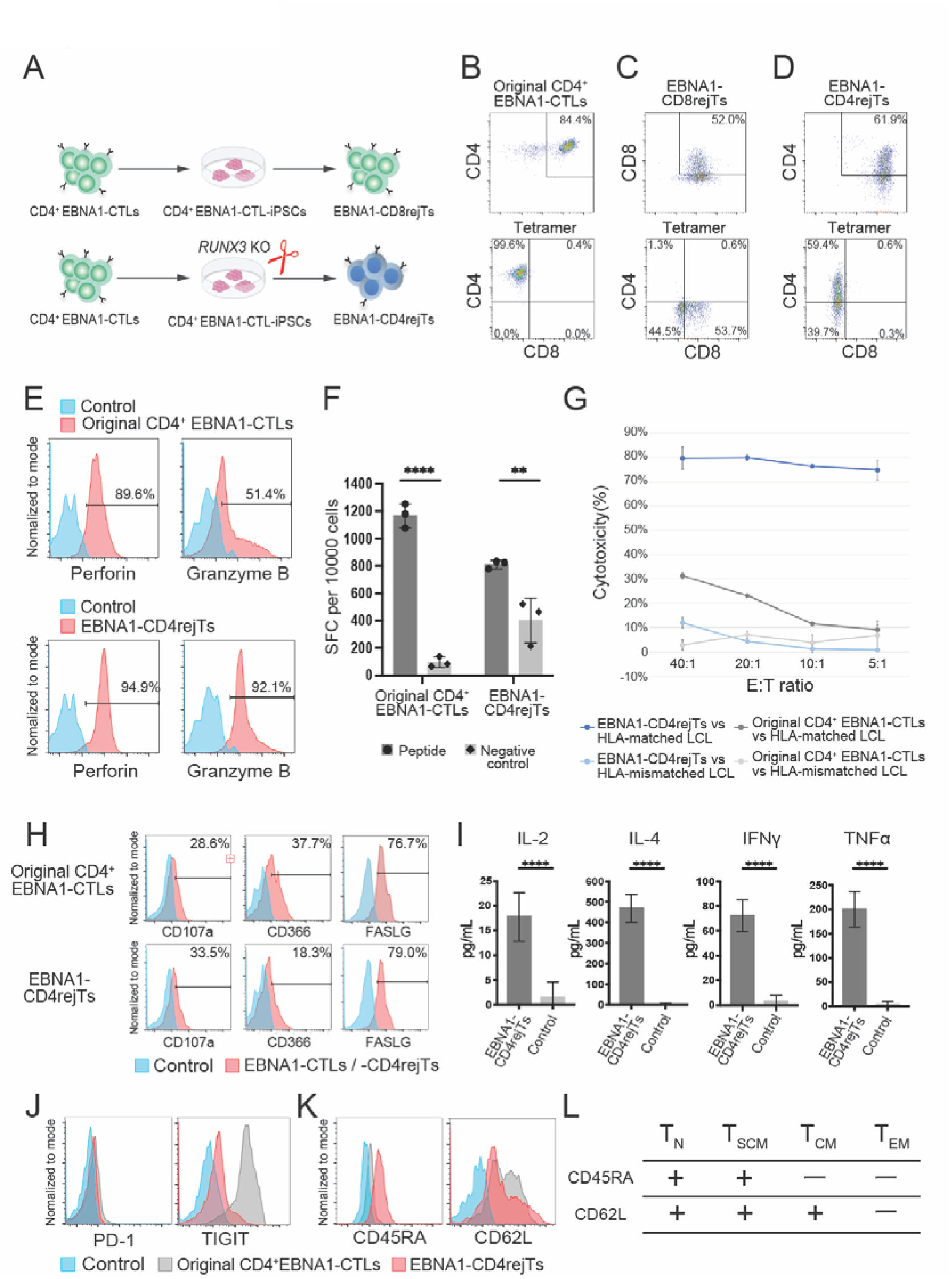
Generation of MHC class II-restricted CD4rejTs from iPSCs (A) Schematic illustration of generation of EBNA1-CD8rejTs (top) and EBNA1-CD4rejTs (bottom). Epstein-Barr nuclear antigen1; EBNA1. (B-D) Flow cytometric CD4/CD8 analysis and tetramer analysis; (B) Original CD4^+^ EBNA1-CTLs, (C) EBNA1-CD8rejTs, (D) EBNA1-CD4rejTs. (E) Histograms of fluorescence intensity expression of intracellular staining perforin and granzyme B for original CD4^+^ EBNA1-CTLs and EBNA1-CD4rejTs (red). Negative controls (fluorescence-minus-one control) are shown in blue. (F) IFN-γ Elispot in response to stimulation with peptide pulse. Negative control shows IFN-γ release in response to no peptide pulse. Results are expressed as spot-forming cells (SFC) per 1.0 x 10^5^ cells. Representative data of at least two independent experiments. Error bars represent ±SD. (G) *In vitro* ^51^Cr assay with EBNA1-CD4rejTs and original CD4^+^ EBNA1-CTLs as effectors against HLA-matched LCLs and HLA-mismatch LCLs as targets. The effector-to-target (E:T) ratios were 40:1, 20:1,10:1, 5:1. Representative data of at least two independent experiments. Error bars represent ±SD. LCL; EBV-infected lymphoblastoid cells. (H) Histograms of fluorescence intensity of expression of surface markers related to Th1 cells (CD107a, CD366, FASLG) for original CD4^+^ EBNA1-CTLs (top) and EBNA1-CD4rejTs (bottom) are shown in red. Negative controls (fluorescence-minus-one controls) are shown in blue. (I) Cytometric bead array for measuring cytokines (high-sensitivity IL-2, IL-4, IFN-γ and TNFα) produced by EBNA1-CD4rejTs after 24 of coculture with NK-YS cells. Error bars represent ± SEM. *p < 0.05, ****p < 0.0001, one-way ANOVA. (J) Histograms of PD-1 and TIGIT expression on EBNA1-CD4rejTs compared with that on original CD4^+^ EBNA1-CTLs and negative controls. (K) Histograms of CD45RA and CD62L expression on EBNA1-CD4rejTs compared with that on original CD4^+^ EBNA1-CTLs and negative controls (fluorescence-minus-one controls). (L) Differention markers characterizing various memory phenotypes. Tn, naïve T cells; Tscm, stem cell memory T cells; Tcm, central memory T cells; Tem, effector memory T cells.

We first generated MHC class II (HLA-DRB1*0101)-restricted CD4^+^ Epstein-Barr nuclear antigen 1 (EBNA1)-specific CTLs (CD4^+^ EBNA1-CTLs) from a healthy donor (Figure 5B) and subsequently reprogrammed them into iPSCs (CD4^+^ EBNA1-CTL-iPSCs). When we differentiated CD4^+^ EBNA1-CTL-iPSCs using our culture platform, we could only generate CD8^+^ EBNA1-specific rejTs (EBNA1-CD8rejTs), as expected (Figure 5C). However, following the knock out of *RUNX3* in these iPSCs, we successfully differentiated them into CD4^+^ EBNA1-specific rejuvenated CTLs (EBNA1-CD4rejTs) instead of CD8rejTs. The antigen specificity against EBNA1 was determined by staining with HLA-DRB1*0101/EBNA1_515-527_ tetramer. Generated EBNA1-CD4rejTs showed almost 100% antigen specificity and 61.9% of these were CD4 SP cells, and none were CD8 positive cells (Figure 5D).

We next assessed the cytotoxic capabilities of EBNA1-CD4rejTs. We evaluated the expression levels of intracellular perforin and granzyme B. EBNA1-CD4rejTs exhibited higher expression levels of both perforin and granzyme B (94.9% and 92.1%, respectively) compared to the original CD4^+^ EBNA1-CTLs (89.6% and 51.4%, respectively) (Figure 5E). IFN-γ levels for original CD4^+^ EBNA1-CTLs and EBNA1-CD4rejTs were measured by IFN-γ ELISPOT assays after stimulation with EBNA1 peptide. Both original CD4^+^ EBNA1-CTLs and EBNA1-CD4rejTs showed specific activity against EBNA1 (1169 ± 88.43 and 827.5 ± 4.95 IFN-γ spot-forming cells [SFCs]/10,000, respectively) (Figure 5F). We also conducted ^51^Cr release assays. The cytotoxicity of EBNA1-CD4rejTs and original CD4^+^ EBNA1-CTLs against HLA-matched and HLA-mismatched EBNA1 peptide-pulsed LCL cells was measured (Figure 5G). At an E:T ratio of 40:1, 20:1, 10:1 and 5:1, the percentage of HLA-matched LCL cells lysed by EBNA1-CD4rejTs was 79.7 ± 4.6%, 80.0 ± 1.2%, 76.4 ± 0.7% and 74.8 ± 4.1%, respectively. Also, at an E:T ratio of 40:1, 20:1, 10:1 and 5:1, the percentage of HLA-matched LCL cells lysed by original CD4^+^ EBNA1-CTLs was 31.2 ± 1.1%, 23.0 ± 0.8%, 11.4 ± 0.7% and 9.0 ± 1.8%, respectively. Taken together, EBNA1-CD4rejTs demonstrated much stronger cytotoxicity with stronger proliferative capability than original CD4^+^ EBNA1-CTLs.

Furthermore, we investigated T cell phenotype of EBNA1-CD4rejTs. EBNA1-CD4rejTs expressed the cell-surface markers CD107a, CD366, and FASLG (33.5%, 18.3%, 79.0%, respectively) at similar levels as original CD4^+^ EBNA1-CTLs (28.6%, 37.7%, 76.7%, respectively) (Figure 5H). We next measured cytokine release levels of EBNA1-CD4rejTs and EBNA1-CD4rejTs secreted IL-2, IL-4, IFN-γ and TNF-α at significant levels compared to negative control (all, p<0.0001) (Figure 5I). To evaluate exhaustion levels, we analyzed PD-1 and TIGIT expression on original CD4^+^ EBNA1-CTLs and EBNA1-CD4rejTs by flow cytometry. The expression of PD-1 on original CD4^+^ EBNA1-CTLs was higher than EBNA1-CD4rejTs (15.1 ± 1.4% vs 4.8 ± 1.6%), whereas that of TIGIT on original CD4^+^ EBNA1-CTLs was much higher than EBNA1-CD4rejTs (94.9 ± 3.3% vs 11.8± 0.9%) (Figure 5J). We also examined memory phenotype of original CD4^+^ EBNA1-CTLs and EBNA1-CD4rejTs. EBNA1-CD4rejTs demonstrated younger memory phenotype (CD45RA^+^, CD62L^+^; T_SCM_) compared with original CD4^+^EBNA1-CTLs (CD45RA^-^, CD62L^+^; T_CM_) (Figure 5K and L)^43^.

Collectively, the generated class II-restricted iPSC-derived CD4^+^ EBNA1-CTLs possess both cytotoxic and helper characteristics.

## Discussion

In this study, we explored the generation of Treg cells from ATL-derived induced iPSCs. Notably, the Treg phenotype CD4^+^ T cells we generated did not express CADM1, a marker commonly used to define ATL cells^18–20^. A stark contrast in gene expression was observed between our generated Treg phenotype CD4^+^ T cells and primary ATL cells, particularly for *KLRB1* (CD161). The CD161 molecule, a C-type lectin-like receptor predominantly expressed on most natural killer (NK) cells and a subset of T cells, carries significant functional implications. CD161^+^ Tregs exhibit a profound suppressive effect, which has been associated with mediating wound healing in autoimmune diseases^35, 36^. This mechanism is largely attributed to the binding of CD161 to its ligand, LLT1, found on T cells in inflamed tissues^37, 38^. Consequently, ATL-derived iPSCs show great promise for not only delineating the underlying pathogenesis of ATL but also for inducing and analyzing Tregs with therapeutic potential against autoimmune conditions, transplant rejection, and graft-versus-host disease.

Our gene expression analysis identified *RUNX3* as a crucial regulator in determining CD4/CD8 lineage choices. It has been documented that the deletion of *Runx3* in a mouse model results in a marked reduction of CD8 single-positive (SP) cells from 95.2% to 33.8%, while CD4 SP cells increase from 0.01% to 2.2%^39^. In human models, *RUNX3* knockout led to differentiation of iPSCs primarily into CD4 SP cells, with minimal generation of CD8 SP cells. These discrepancies may stem from species differences or potential *in vivo* redundancies in the transgenic mouse models.

Conversely, the overexpression of *RUNX3* was able to induce the formation of CD8 SP cells even from ATL-derived iPSCs. This finding strongly supports the notion that *RUNX3* is a pivotal regulator of T cell lineage commitment. Furthermore, our weekly RNA sequencing analyses of EBV-CD8rejTs and EBV-CD4rejTs throughout T cell differentiation revealed that *RUNX3* knockout in iPSCs downregulated genes associated with CD8 lineage commitment (such as *GIMAP5*, *ITGAE*, *CST7*), while genes linked to CD4 lineage (like *ZBTB7B*, *CD40LG*) remained relatively stable (Supplementary Figure 4). Although *GIMAP5*, *ITGAE*, and *CST7* emerged as candidate genes in generating CD4 SP cells, the significant downregulation of these genes in *RUNX3* knockout further confirms the upstream role of *RUNX3* in T cell lineage determination.

Interestingly, our approach of knocking out *RUNX3* in iPSCs derived from antigen-specific CD8^+^ CTLs successfully led to the generation of MHC class I-restricted CD4 SP Th1 cells. While the presence of such cells has been noted in previous studies, they remain relatively scarce^40, 41^. Our *in vitro* differentiation system from iPSCs derived from mature T cells allows for the generation of a diverse array of T cells, effectively bypassing the selection processes typically occurring in the thymus.

Th1 cells are recognized for their ability to counteract CAR-T cell exhaustion, thereby enhancing persistence and efficacy in antitumor responses^15^. For instance, Lisocabtagene Maraleucel, a second-generation CAR-T therapy targeting CD19, has shown promising clinical outcomes in relapsed or refractory lymphomas when administered with equal doses of CD8^+^ and CD4^+^ CAR-T cells at a 1:1 ratio^42^. The Th1 phenotype EBV-CD4rejTs generated in our study displayed functional characteristics that enhanced the cytotoxicity of CD8^+^ T cells, both *in vitro* and *in vivo* (Figure 4).

Moreover, through *RUNX3* knockout in iPSCs derived from CD4^+^ EBNA1-CTLs, we successfully generated MHC class II-restricted EBNA1-CD4rejTs, which exhibited both cytotoxic and helper functionalities. These EBNA1-CD4rejTs expressed less TIGIT expression and showed younger memory phenotype (stem cell memory phenotype)^43^ than original CD4^+^ EBNA1-CTLs (Figure 5). This innovative strategy enables the production of an unlimited supply of iPSC-derived CD4^+^ CTLs, a crucial component for effective immune responses against viral infections and cancer^33, 34^.

By selectively knocking out *RUNX3* in iPSCs, we are now able to generate an unlimited number of therapeutic CD4^+^ T cells that enhance cytotoxicity, mitigate exhaustion in CD8^+^ CAR-T cells, and promote the development of memory phenotype cells. We have successfully established iPSCs derived from healthy donor CTLs and edited their HLA class I to facilitate allogeneic adoptive T cell therapy targeting cervical cancer. We have validated these cells and started an investigator-initiated clinical trial. The method described in this study for generating CD4^+^ T cells will enable the rapid production of Th1 cells, thereby enhancing the efficacy of T cell therapies.

In conclusion, our method of generating CD4 SP cells from iPSCs represents a straightforward yet promising approach for developing “off-the-shelf” immunotherapies that leverage the helper and cytotoxic functions of CD4^+^ Th1 or CD4^+^ CTLs. This technique has the potential to significantly advance therapeutic strategies in the fight against cancer, viral infection and autoimmune disorders, thus opening new avenues for effective immunotherapy.

## Materials and methods

### Cell lines and cell culture

We recruited donors through the Department of Hematology, Juntendo University School of Medicine. This study was conducted in accordance with the Declaration of Helsinki and the Ethics Committee at Juntendo University School of Medicine approved the experimental protocol. Peripheral blood from two ATL patient (Supplementary Table 1) and healthy donors was obtained with informed consent. Peripheral blood was sorted using Moflo Astrios EQs (Beckman Coulter, Brea, CA) and a BD FACSAria III cell sorter (BD Bioscience, San Jose, CA). ATL cells were transduced with Sendai virus vectors encoding the Yamanaka 4 factors (OCT3/4, SOX2, KLF4, and c-MYC) and SV40 large T antigen. Healthy donor-derived CTL clones were transduced with Sendai virus vector encoding the Yamanaka 4 factors with NANOG and LIN28 ^3, 7^. Transduced cells were seeded onto plates coated with iMatrix-511 (Nippi, Tokyo, Japan). Seeded cells were cultured in NS-A2 (Shimadzu Diagnostic, Tokyo, Japan) medium in the presence of 100 U/mL of IL-2 (Miltenyi Biotec, Bergisch Gladbach, Germany), which was later replaced with StemFit AK03N (Ajinomoto Healthy Supply, Tokyo, Japan). Human healthy donor CTL cell lines (CD4^+^ T cell clone; HLA-A*2402-restricted LMP2_419-427_-specific CD8^+^ CTLs; HLA-A*2402-restricted HPV 16 E6_49-57_-specific CD8^+^ CTLs; HLA-DRB1*0101-restricted EBNA1_515-527_-specific CD4^+^ CTLs) and human iPSC lines (CD4-iPSCs; EBV-iPSCs; HPV-iPSCs; EBNA1-CTL-derived iPSCs) were established as described ^2, 3^.

Extranodal natural killer/T-cell lymphoma, nasal type, cell line NK-YS was a gift from Dr. J. Tsuchiyama (Okayama University Medical School, Okayama, Japan). NK-YS cells were grown in Iscove’s modified Dulbecco’s medium (IMDM, Thermo Fisher Scientific, Waltham, MA) supplemented with 10% FBS (FBS; Gibco Life Technologies, Carlsbad, CA), 1% penicillin-streptomycin-glutamine (PSG; Thermo Fisher Scientific), and 100 IU/mL IL-2 (Miltenyi Biotec). Extranodal natural killer/T-cell lymphoma, nasal type, ENKL-J1 cells ^44^ and CAEBV cells ^45^ were established and cultured as described. The HLA-A24^+^ cervical cancer line SiHa was purchased from American Type Culture Collection (ATCC; Manassas, VA) and the cervical cancer cell line SKG-IIIa was purchased from RIKEN BioResource Research Center (RIKEN BRC; Ibaragi, Japan). Cervical cancer cell lines were cultured in Rosewell Park Memorial Institute (RPMI) 1640 medium (Sigma-Aldrich, Saint Louis, MO) supplemented with 10% FBS (Gibco Life Technologies) and 1% PSG (Thermo Fischer Scientific).

### Reprogramming sorted T cells into iPSCs

ATL cells were transduced with Sendai virus vectors encoding the Yamanaka 4 factors (OCT3/4, SOX2, KLF4, and c-MYC) and SV40 large T antigen. Healthy donor-derived CTL clones were transduced with Sendai virus vector encoding the Yamanaka 4 factors with NANOG and LIN28 ^3, 7^. Transduced cells were seeded onto plates coated with iMatrix-511 (Nippi). Seeded cells were cultured in NS-A2 medium (Shimadzu Diagnostic) in the presence of 100 U/mL of IL-2 (Miltenyi Biotec), which was later replaced with StemFit AK03N (Ajinomoto Healthy Supply).

### Differentiation of iPSCs

iPSCs were differentiated into rejTs as described ^1, 16, 17^. Briefly, we manually picked up each iPSC colony when stable colonies were found. These small clumps of iPSCs were transferred onto C3H10T1/2 cells with coculture in Iscove’s modified Dulbecco’s medium (Sigma-Aldrich) supplemented with 15% fetal bovine serum (FBS) (HyClone, GE Healthcare, South Logan, UT) and a cocktail of 10 mg/mL human insulin, 5.5 mg/mL human transferrin, 5 ng/mL sodium selenite, 2 mM L-glutamine (all Thermo Fisher Scientific), 0.45 mM α-monothioglycerol (Wako Pure Chemicals, Osaka, Japan), and 50 mg/mL vascular endothelial growth factor (Miltenyi Biotec).

On day 14 of coculture, sac-like structures containing hematopoietic progenitor cells were extracted and transferred onto DL1/4-expressing C3H10T1/2 feeder cells for T cell lineage differentiation during coculture in α-minimum essential medium (Thermo Fisher Scientific) supplemented with 20% FBS (HyClone), as well as penicillin-streptomycin-glutamine (PSG) in the presence of 20 ng/mL recombinant human stem cell factor, 10 ng/mL Fms-related tyrosine kinase 3 ligand, and 10 ng/mL IL-7 (all Miltenyi Biotec). On day 28 of coculture, T-lineage cells were harvested and stimulated with 5 mg/mL phytohemagglutinin-L (Sigma-Aldrich), mixed with irradiated peripheral blood mononuclear cells (PBMCs), and cocultured in CTL medium in the presence of 10 ng/mL IL-2 (Miltenyi Biotec) for CD4^+^ T cell generation and 10 ng/mL IL-7 and IL-15 (both, Miltenyi Biotec) for CD8^+^ T cell generation. Media such as C3H10T1/2, the DL1/4-expressing C3H10T1/2 feeder cell lines, and gamma-irradiated FBS (HyClone) have already been validated and are compatible with good manufacturing practice standards for clinical use.

### T cell expansion

T cells (1.0 x 10^6^ cells/well) were plated on 24-well culture plates with 1mL NS-A2 (Shimadzu Diagnostic) supplemented with 1% PSG. Every 2 weeks, cells were counted and stimulated with PHA-L (5 µg/mL) mixed with 2.5 x 10^6^/mL of irradiated PBMCs in the presence of IL-2 (Miltenyi Biotec) (10ng/mL) or IL-7 and IL-15 (both Miltenyi Biotec, 10ng/mL). CD4 SP cells and CD8 SP cells were sorted using CD4 Micro Beads and CD8 Micro Beads magnetic cell separation (Miltenyi Biotec).

### Antibodies

All antibodies used in this study were chromophore-conjugated monoclonal antibodies.

To confirm surface antigens, allophycocyanin (APC)/cyanin 7 (Cy7)-conjugated mouse anti-human CD3, APC-conjugated mouse anti-human CD4, APC-conjugated mouse anti-human CCR4 (CD194), Alexa Fluor 750-conjugated mouse anti-human CD45RA, Pacific Blue-conjugated mouse anti-human CD45RA, phycoerythrin (PE)/Cy7-conjugated mouse anti-human CD62L, PE/Cy7-conjugated mouse anti-human CD8, PE-conjugated mouse anti-human CD4 (all BioLegend, San Diego, CA), APC-conjugated mouse anti-human CD7, Brilliant Violet 421-conjugated mouse anti-human CD4, fluorescein isothiocyanate (FITC)-conjugated mouse anti-human CD8, FITC-conjugated mouse anti-human CD3, PE/Cy7-conjugated mouse anti-human CD25, V500-conjugated mouse anti-human CD8, V500-conjugated mouse anti-human CD45 (all BD Bioscience, San Jose, CA), Alexa Fluor 700-conjugated mouse anti-human CD3 (eBioscience, San Diego, CA) and PE-conjugated chicken anti-human TSLC/CADM1 (MBL International, Nagoya, Japan) were used.

For intracellular staining of FOXP3 we used PE-conjugated mouse anti-human FOXP3 (BioLegend). For intracellular staining of perforin and granzyme B we used PE/Cy7 anti-human perforin (BioLegend) and Alexa 647 anti-human granzyme B (BioLegend).

To examine expression of exhaustion markers, BV 605 anti-human CD223 (LAG-3) antibody, APC mouse anti-human programmed death-1 (PD-1), and APC/Cy7 anti-human CD366 (Tim-3) antibody (all BioLegend) were used.

### Flow cytometry

Flow cytometry used BD LSRFortessa equipment (BD Biosciences). The results were analyzed with FlowJo software 10.5.3 (Tree Star, Ashland, OR).

Fluorescence-minus-one controls were used in all antibody combinations to interpret flow cytometry data. The cell suspensions were incubated with antibody cocktails at 4℃ in the dark for 30 min. Afterward, the cells were washed once with 6 mL PBS. For intracellular staining, eBioscience Foxp3/Transcription Factor staining Buffer Set was used according to manufacturer protocol (Thermo Fisher Scientific).

### Quantitative real-time PCR

Total RNA was extracted from established iPSCs and rejTs using TRIzol (Invitrogen) and cDNA was synthesized using a ReverTra Ace qPCR RT kit (Toyobo, Osaka, Japan). All quantitative real-time PCR procedures were carried out on the StepOnePlus real-time PCR system (Applied Biosystem, Carlsbad, CA). Individual PCR reactions were normalized against *GAPDH* levels.

### 51Cr release assay

The cytotoxicity of effector cells against target cells, as analyzed by ^51^Cr release assay, was determined at different E:T ratios ^46^. Target cells were labeled with ^51^Cr (PerkinElmer, Waltham, MA) for 1 h at 37℃ and effector cells were cocultured with the target cells for 6 h. Some target cells were also pulsed with synthesized peptides (50 ng) at 37℃ for 30 min. After 6 h of incubation, supernatant was harvested and radioactivity was measured with a 1450 MicroBeta TriLux scintillation counter (PerkinElmer). Cytolysis was calculated as ([experimental release − spontaneous release]/[maximum release − spontaneous release]) × 100 (%).

To investigate the function of ATL-CD4rejTs, the cytotoxicity of CTL specific to HTLV-1 viral oncoprotein antigen Tax (Tax-CTL) against primary ATL cells with or without the addition of ATL-CD4rejTs was evaluated at E:T ratios of 10:1 and 5:1. ATL-CD4rejTs were mixed with an equal number of Tax-CTLs.

The cytotoxicity of EBV-CD8rejTs and EBV-CD4rejTs against HLA-A*2402^+^ NK-YS ^47^, ENKL-J1 ^44^, and CAEBV cells ^45^ was measured by a ^51^Cr release assay at E:T ratios of 20:1 and 10:1. Target cells were pulsed with HLA-A*2402/LMP2_419-427_ custom-synthesized peptides (Mimotopes, Mulgrave, VIC, Australia). The cytotoxicity of HPV-CD8rejTs and HPV-CD4rejTs against SKGIIIa and SiHa cells was measured by ^51^Cr release assay at E:T ratios from 40:1 to 5:1. Target cells were pulsed with HLA-A*2402/HPV16E6_419-427_ peptide (Mimotopes). The cytotoxicity of GD2-CARTs cocultured with EBV-CD4rejTs against ENKL-J1 was measured by ^51^Cr release assay at E:T ratios from 40:1 to 5:1. Target cells were pulsed with HLA-A*2402/LMP2_419-427_ peptide (Mimotopes). The cytotoxicity of original CD4^+^ EBNA1-CTLs and EBNA1-CD4rejTs against LCL cells was measured by ^51^Cr release assay at E:T ratios from 40:1 to 5:1. Target cells were pulsed with HLA-DRB1*0101/EBNA1_515-527_ peptide (Mimotopes).

Synergy of drug combination used for ^51^Cr release assay was determined using the highest-single-agent model of BIGL 1.4.3 package / Seurat R package (version 4.3.0) as described ^48, 49^.

### Cellular indexing of transcriptomes and epitopes by sequencing (CITE-seq), single-cell RNA sequencing, and single-cell data processing

CITE-seq was performed by staining and barcoding antibodies to cells according to the manufacturer’s protocol ^50^. Briefly, approximately 2.0 x10^6^ cells / sample with viability >85% were suspended in phosphate-buffered saline (PBS) + 0.04% bovine serum albumin (BSA) (Thermo Fisher Scientific) and incubated for 10 min with 5μl of TruStain FcX blocking antibodies (BioLegend). Subsequently, cells were incubated with TotalSeq^TM^-B Human Universal Cocktail, V1.0 (Biolegend) for 30 min at 4°C. After staining, cells were washed 4x (once with PBS+ 0.04% BSA, then three times with PBS+ 1% BSA), followed by centrifugation (400 rpm 5min at 4°C) and supernatant exchange. Cells were then processed for single-cell RNA sequencing using the 10× Chromium system with the Chromium Next Gen Single Cell 3’ Reagent Kits v 3.1 (Dual Index) (10× Genomics, Pleasanton, CA). Bio-Rad T100 thermal cycler (Bio-Rad Laboratories, Hercules, CA) was used for PCR. After the libraries were qualified using Bioanalyzer 2100 (Agilent, Tokyo, Japan), samples were subjected to sequencing with HiseqX (Illumina, San Diego, CA). Raw reads (fastq files) were re-aligned using Cell Ranger (v7.0.0) (10× Genomics). Normalization and downstream analysis of RNA data were performed using the Seurat R package (version 4.3.0) ^49^, which enables the integrated processing of multi-modal single cell datasets, and Python v3.13^51^, which enables the integrated processing of multi-modal single cell datasets. We used CellTypist automated cell type annotation for scRNA-seq datasets ^32^. Automated cell type annotation was performed according to expression of various gene sets, including those for Th1 cells (*TBX21*, *CXCR3*, *KLRB1*, *DUSP2*, *RGCC*, *JUND*, *CEBPB*, *DNAJA4*, *etc*.), Tcm/Naive cytotoxic T cells (*YBX3*, *CD8B*, *HLA-A*, *S100B*, *KLRB1*, *B2M*, *HLA-C*, *etc*.), Tcm/Trm cytotoxic T cells (*CMC1*, *CST7*, *SH2D1A*, *etc*.), Tcm/Naive helper T cells (*MT-RNR2*, *SELL*, *IGFBP1*, *FCGRT*, *TSHZ2*, *HBG1*, *RPL17*, *EEF1G*, *etc*.), MAIT cells (*IL7R*, *KLRG1*, *SLC4A10*, *TRAV1-2*, *NCR3*, *etc*.), and large pre-B cells (*MME*, *CD24*, *MKI67*, *IGLL1*, etc.) ^32^.

### CRISPR/Cas9 gene editing

iPSCs were treated with RevitaCell (Thermo Fisher Scientific) 24 h before electroporation. Cells at 80% confluence were harvested with Accutase (Life Technologies, Coralville, IA). Before electroporation, a nucleofection solution was made by first mixing 150 mg/mL of SpCas9 (Integrated DNA Technologies) and 87.5 mg/mL of sgRNA at 1:3 molar ratio directly, incubating for 10 min at room temperature, then diluting with 20 uL of P3 Primary Cell solution (Lonza, Basel, Switzerland). For each reaction, 500,000 cells were mixed with the nucleofection solution. Nucleofection was performed using 16-well Nucleocuvette Strip with 4D Nucleofector system (Lonza) using CA137 electroporation code. Immediately after electroporation, cells were transferred into one well of an iMatrix-coated 6-well plate containing 500 ml of StemFit medium (Ajinomoto Healthy Supply) with 10 mM Y-27632 (Fujifilm Wako Pure Chemical, Osaka, Japan). Donor vector was added at 100K multiplicity of infection (MOI) (measured by qPCR as described) directly to cells after plating and incubated at 37LJ for 24 h. Medium was changed 24 h after editing and 10 mM Y-27632 was removed 48 h after edit. Synthetic sgRNAs were purchased from Synthego (Redwood City, CA). The genomic sgRNA target sequence for CD4-iPSCs was CACUGCGGCCCACGAAGCGA and those for sgRNA *RUNX3* were GGTGGTGACTGTGATGGCAGGC (forward) and AGAGGGGGTGGCATGAACGGTTTC (reverse).

### RNA extraction, quantification, and sequencing

RNA was extracted using the RNeasy Micro Kit (QIAGEN, Hilden, Germany), then quantified using a NanoDrop 2000 (Thermo Fisher Scientific) according to manufacturer instructions. For RNA sample preparations, 10 ng of RNA per sample was used as the input material. Libraries were generated using a SMART-Seq v4 Ultra Low Input RNA Kit (Takara Bio USA, Mountain View, CA, USA) following the manufacturer’s recommendations.

### Interferon-**γ** (IFN-**γ**) enzyme-linked immunospot (ELISPOT) assays

IFN-γ ELISPOT assays were performed as described ^52, 53^. Briefly, 96-well filter plates (Merck Millipore, Darmstadt, Germany) were coated with 10 mg/mL of an anti-IFN-γ monoclonal antibody (mAb), 1-DIK (Mabtech, Stockholm, Sweden). 5.0 x10^4^ cells were added to the wells in the presence of peptides at a final concentration of 10 ug/ml for each peptide. The plates were incubated overnight at 37LJ in 5% CO2. The cells were discarded the following day and a biotinylated anti-IFN-γ mAb, 7-B6-1 (Mabtech), was added at 1 mg/mL and left for 2 h at 37LJ, followed by the addition of streptavidin-horseradish peroxidase (Mabtech) for 1 h at room temperature. Spot color was developed by adding AP conjugate substrate kit (Bio-Rad Laboratories). Plate color was quantified using Elispot classic reader AID-ELR08-1SYSTEM (AID, Strassberg, Germany). The results are shown as the number of spot-forming cells (SFC)/1.0 x10^4^ cells.

### Analysis of cytokine release capacity through cytometric bead array

To determine the concentrations of the cytokines (high-sensitivity IL-2, IL-4, IL-10, TNF-α, and IFN-γ) released into culture media from EBV-CD8rejTs, EBV-CD4rejTs and EBNA1-CD4rejTs, the BD Cytometric Bead Array (CBA) Human Th1/Th2/Th17 Cytokine Kit (BD Biosciences) was used according to manufacturer’s instructions. 1.0 × 10^5^ cells were cultured in 96-well plates for 16 h, with collection of supernatant samples. Levels of multiple cytokines were simultaneously determined by flow cytometry (LSRFortessa, BD Biosciences). The obtained data were analyzed using CBA software (BD Biosciences).

### Generation of peripheral blood-derived CART cells

Healthy donor PBMC were activated by T cell Transact (Miltenyi Biotec) in the presence of 100 U/mL IL-2 (Miltenyi Biotec). Three days later, activated T cells were plated on 24-well plates coated with RetroNectin (Takara Bio) and retroviral GD2-CAR supernatant was added to each well. GD2-CART cells were cultured in NS-A2 (Nissui, Tokyo, Japan) with 10 ng/mL each of IL-7 and IL-15 (both Miltenyi Biotec). CAR-positive T cells were sorted so that CAR transgene expression in GD2-CART cells was approximately 80%.

### Coculture experiment and examination of exhaustion markers and memory phenotype

GD2-CARTs, EBV-CD4rejTs, and GD2-CARTs mixed with EBV-CD4rejTs were cocultured with ENKL-J1 cells at a 1:4 ratio (24000 cells : 96000 cells) ^54^. All samples’ viability was > 80%. Flow cytometry was performed on day 7 of coculture to examine exhaustion markers (TIGIT, TIM-3, LAG-3, and PD-1) and memory phenotype. We defined memory phenotype subsets based on expression of the cell-surface markers CD45RA, CD62L, CD95, and CD28. We defined ENKL-J1 cells as CD3^-^ cells, GD2-CARTs as CD3^+^ CD8^+^ cells, and EBV-CD4rejTs as CD3^+^ CD4^+^ cells.

### *In vivo* experiments

All *in vivo* studies were approved by the Animal Research Committees of Juntendo University School of Medicine. ENKL-J1 cells transduced with γ-retroviral vector encoding *GFP/FFluc* were sorted for GFP expression by flow cytometry. Six-week-old female NOD/Shi-scid, IL-2RγKO Jic (NOG) mice (In-Vivo Science, Tokyo, Japan) were intraperitoneally engrafted with *GFP/FFluc* ENKL-J1 (1.0 x10^6^ cells). Mice were divided into 4 groups (3 treatment groups and one untreated group). Five days after tumor inoculation, mice in treatment groups were intraperitoneally injected with CD8^+^/CAR^+^ sorted GD2-CARTs, EBV-CD4rejTs, or CD8^+^/CAR^+^ sorted GD2-CARTs mixed with EBV-CD4rejTs (4.0×10^6^ cells, one dose on Day 0) to permit observation of the anti-tumor effect in each group. Coculture of CD8^+^/CAR^+^ sorted GD2-CARTs and EBV-CD4rejTs at a 1:1 ratio, without cytokines, began 3 days before treatment.

Tumor burden was monitored using the Xenogen IVIS imaging system (Xenogen, Alameda, CA). Firefly D-luciferin substrate (OZ Biosciences, Marseille, France) was intraperitoneally injected into mice 15 min before imaging. Living Image software version 4.7.2 (PerkinElmer) was used for luminescence analyses. Intensity signals were measured as total photon/s/cm^2^/steradian (p/s/cm^2^/sr).

### Statistical analysis

All data are presented as means ± standard deviation (SD). Results were analyzed using unpaired Student’s *t*-test (two-tailed) or ordinary one-way or two-way analysis of variance as stated, with a *p* value <0.05 indicating a significant difference. All data are presented as mean ± SD as stated in the Figure legends. Software used for statistical analyses was Excel (Microsoft, Redmond, WA) and Prism 8.9 (GraphPad Software, San Diego, CA).

---

Supplementary Figure 1. ATL-CD4rejTs are CD161^+^Tregs

(A) Flow cytometric analysis of ATL-CD4 rejTs from a different acute type ATL patient. (B) Quantitative real-time PCR analysis to evaluate the relative expression of Tax and HBZ against a positive control (primary ATL cells) in ATL-iPSCs and ATL-CD4rejTs. (C) UMAP of scRNA-seq data showing primary ATL cells (left) and ATL-CD4rejTs (right). Cluster gene characteristics determine cluster functional description. Cluster 0: *FOXP3*^+^ CD4 T cells, cluster 1: *KLRB1*^+^ CD4 T cells, cluster 2:*CADM1*^+^ CD4 T cells, cluster 3: *MKI67*^+^ CD4 T cells, cluster 4: *NKG7*^+^ CD4 T cells and CD8 T cells, cluster 5: *CLNT*^+^ CD4 T cells No. 1, cluster 6: *CLNT*^+^ CD4 T cells No. 2, cluster 7: *ZNF683*^+^ CD4 T cells, cluster 8: *TSHZ2*^+^ CD4 T cells. (D) CITE-seq data of CD4 and CD8A for primary ATL cells and ATL-CD4rejTs. (E) Volcano plots for primary ATL cells (left) and ATL-CD4rejTs (right) showing the most significant differentially-expressed genes (DEG) in each sample. (F) UMAP of *KLRB1*, the most significant DEG in ATL-CD4rejTs.

Supplementary Figure 2. Analysis of genes associated with CD4/CD8 T cell lineage choice

(A) CITE-seq data of CD4 and CD8A for CD8rejTs and ATL-CD4rejTs. (B) UMAPs of genes associated with CD4/CD8 lineage choice in CD8rejTs and ATL-CD4rejTs.

Supplementary Figure 3.Analysis of HPV-rejTs and comparison of EBV-CD4rejTs with conventional T cells

(A) Flow cytometric CD4/CD8 analysis of post-selected HPV-CD8rejTs and HPV-CD4rejTs. (B) *In vitro* ^51^Cr assay with HPV-CD8rejTs and HPV-CD4rejTs as effectors against SKG-IIIa cells (left) and SiHA cells (right). The effector-to-target (E:T) ratios were 40:1, 20:1, 10:1 and 5:1. Representative data of at least three independent experiments. Error bars represent ±SD. (C) Histograms of fluorescence intensity of expression of intracellular staining of perforin and granzyme B for HPV-CD8rejTs and HPV-CD4rejTs (red). Negative controls (fluorescence-minus-one control) are shown in blue. (D) Histograms of fluorescence intensity of expression of surface markers related to Th1 cells (CD107a, CD366, FASLG) for EBV-CD8rejTs and EBV-CD4rejTs (red). Negative controls (fluorescence-minus-one controls) are shown in blue. (E) UMAP analysis of conventional T cells (control T cells) shown in blue and EBV-CD4rejTs shown in orange. (F) Annotation of scRNA-seq data of control T cells and EBV-CD4rejTs using CellTypist automated cell type annotation for scRNA-seq datasets. (G) Annotation of scRNA-seq data of control T cells and EBV-CD4rejTs. Cluster 0: Th1 cells No. 1, cluster 1: MAIT cells, cluster 2: Tcm/Naive cytotoxic T cells, cluster 3: Th1 cells No. 2, cluster 4: Tcm/Naive helper T cells Tcm, cluster 5: large pre-B cells, cluster 6: Tem/Trm cytotoxic T cells. Tcm, central memory T cells; Tem, effector memory T cells; Trm, tissue memory T cells.

Supplementary Figure 4. Weekly RNA-seq data of EBV-CD8rejTs and EBV-CD4rejTs during T cell differentiation Weekly RNA-seq analysis of genes associated with CD4/CD8 lineage choice in EBV-CD8rejTs and EBV-CD4rejTs during T cell differentiation. RNA-seq was performed after collection of iPSC-derived hematopoietic progenitor cells (Day 0) every week until day 28.

## Acknowledgements

We thank A.S. Knisely for critical reading of the manuscript. We thank Tamami Sakanishi for fluorescence-activated cell sorter operation and members of the Laboratory of Radioisotope Research, Research Support Center, Juntendo University Graduate School of Medicine for technical assistance. We thank Satoshi Yamazaki and Takaharu Kimura for helping with scRNA-seq. These studies were supported by a grant from the Japan Society for the Promotion of Science Grants-in-Aid for Scientific Research (KAKENHI) program (Grant Number 24K02330) and a grant from Juntendo University School of Medicine Center for Genomic and Regenerative Medicine (AM425G2309)

## Authorship contributions

Y.F. performed experiments, analyzed the data, and wrote the manuscript. M.I. and S.K performed experiments and analyzed the data. A.G helped with cell culture and *in vivo* experiments. K.M helped perform *in vivo* experiments. K.Y. and T.T. helped with flow cytometry experiments. N.I and K.T. performed scRNA-seq analysis. J.A. helped with experiments and provided scientific discussion. H.N. provided scientific discussion and wrote the manuscript. M.A. planned and directed the study and wrote the manuscript.

