## Supplementary Figures and Tables for "Generation of iPSC-derived CD4^+^ Th1 cells enhancing chimeric antigen receptor-T cell cytotoxicity"

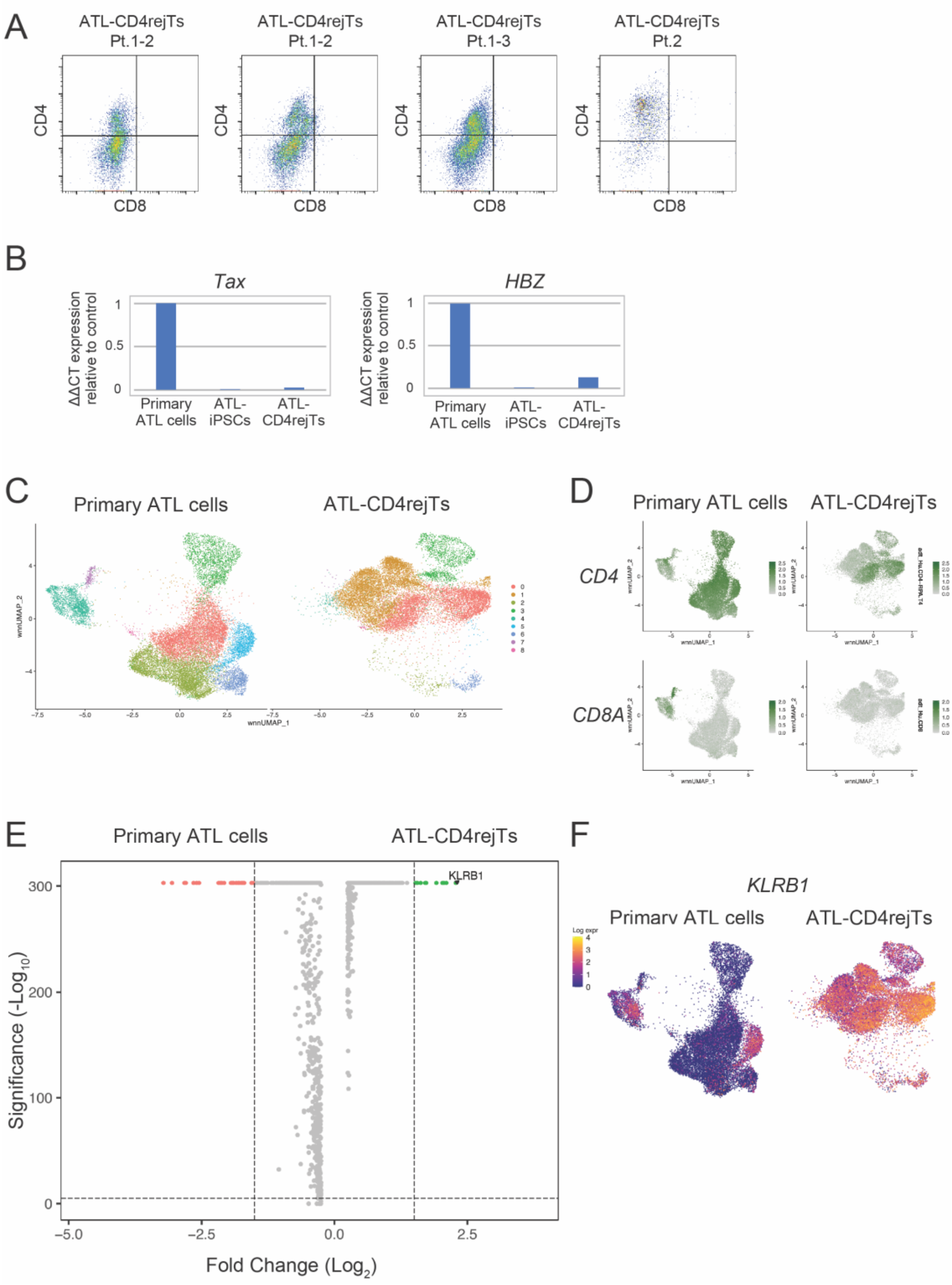


**Supplementary Figure 1. ATL-CD4rejTs are CD161$^{+}$Tregs**

(A) Flow cytometric analysis of ATL-CD4 rejTs from a different acute type ATL patient.

(B) Quantitative real-time PCR analysis to evaluate the relative expression of Tax and HBZ against a positive control (primary ATL cells) in ATL-iPSCs and ATL-CD4rejTs. (C) UMAP of scRNA-seq data showing primary ATL cells (left) and ATL-CD4rejTs (right). Cluster gene characteristics determine cluster functional description. Cluster 0: *FOXP3*$^{+}$ CD4 T cells, cluster 1: *KLRB1*$^{+}$ CD4 T cells, cluster 2:*CADM1*$^{+}$ CD4 T cells, cluster 3: *MKI67*$^{+}$ CD4 T cells, cluster 4: *NKG7*$^{+}$ CD4 T cells and CD8 T cells, cluster 5: *CLNT*$^{+}$ CD4 T cells No. 1, cluster 6: *CLNT*$^{+}$ CD4 T cells No. 2, cluster 7: *ZNF683*$^{+}$ CD4 T cells, cluster 8: *TSHZ2*$^{+}$ CD4 T cells. (D) CITE-seq data of CD4 and CD8A for primary ATL cells and ATL-CD4rejTs. (E) Volcano plots for primary ATL cells (left) and ATL-CD4rejTs (right) showing the most significant differentially-expressed genes (DEG) in each sample. (F) UMAP of *KLRB1*, the most significant DEG in ATL-CD4rejTs.

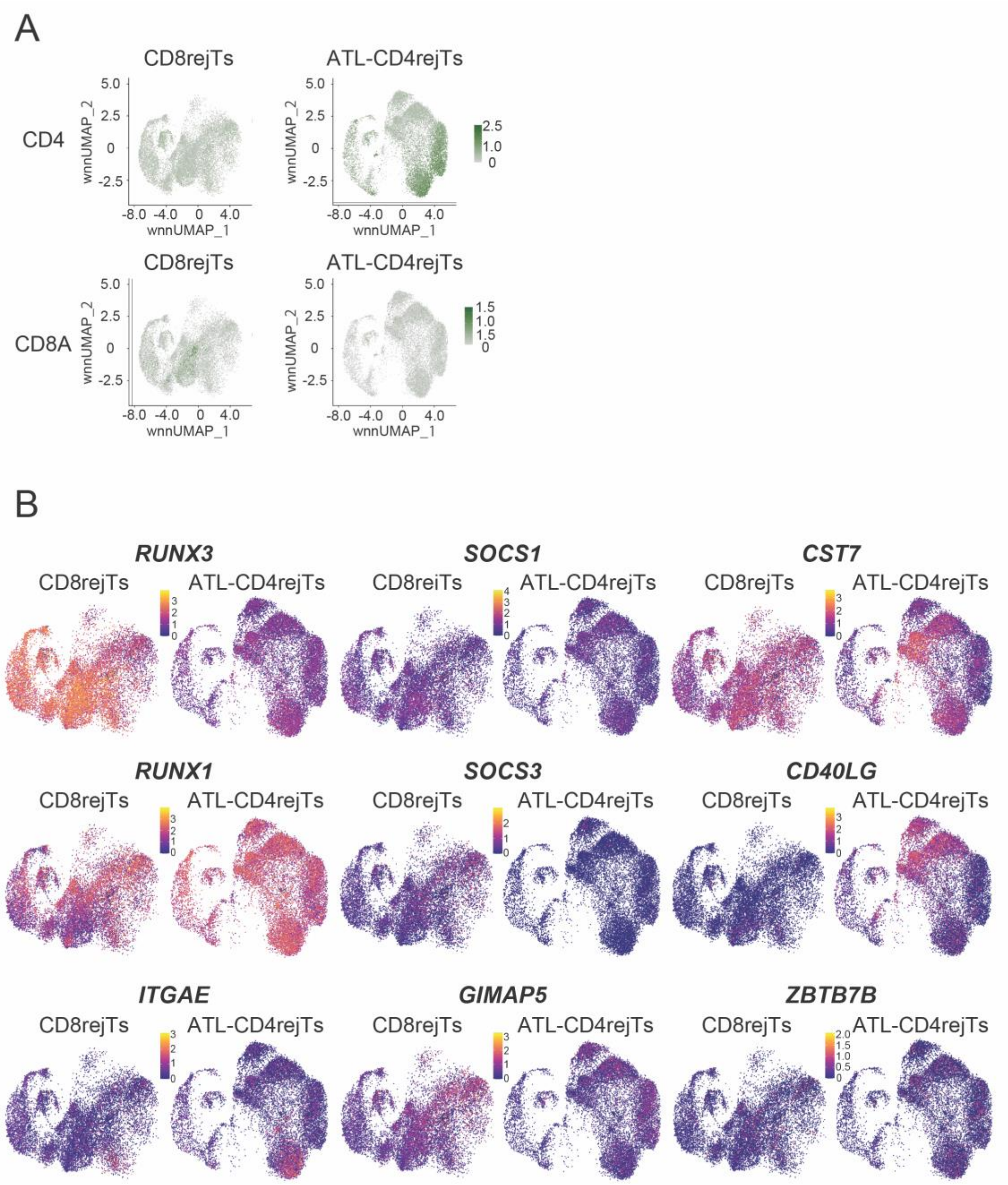


**Supplementary Figure 2. Analysis of genes associated with CD4/CD8 T cell lineage choice**

(A) CITE-seq data of CD4 and CD8A for CD8rejTs and ATL-CD4rejTs. (B) UMAPs of genes associated with CD4/CD8 lineage choice in CD8rejTs and ATL-CD4rejTs.

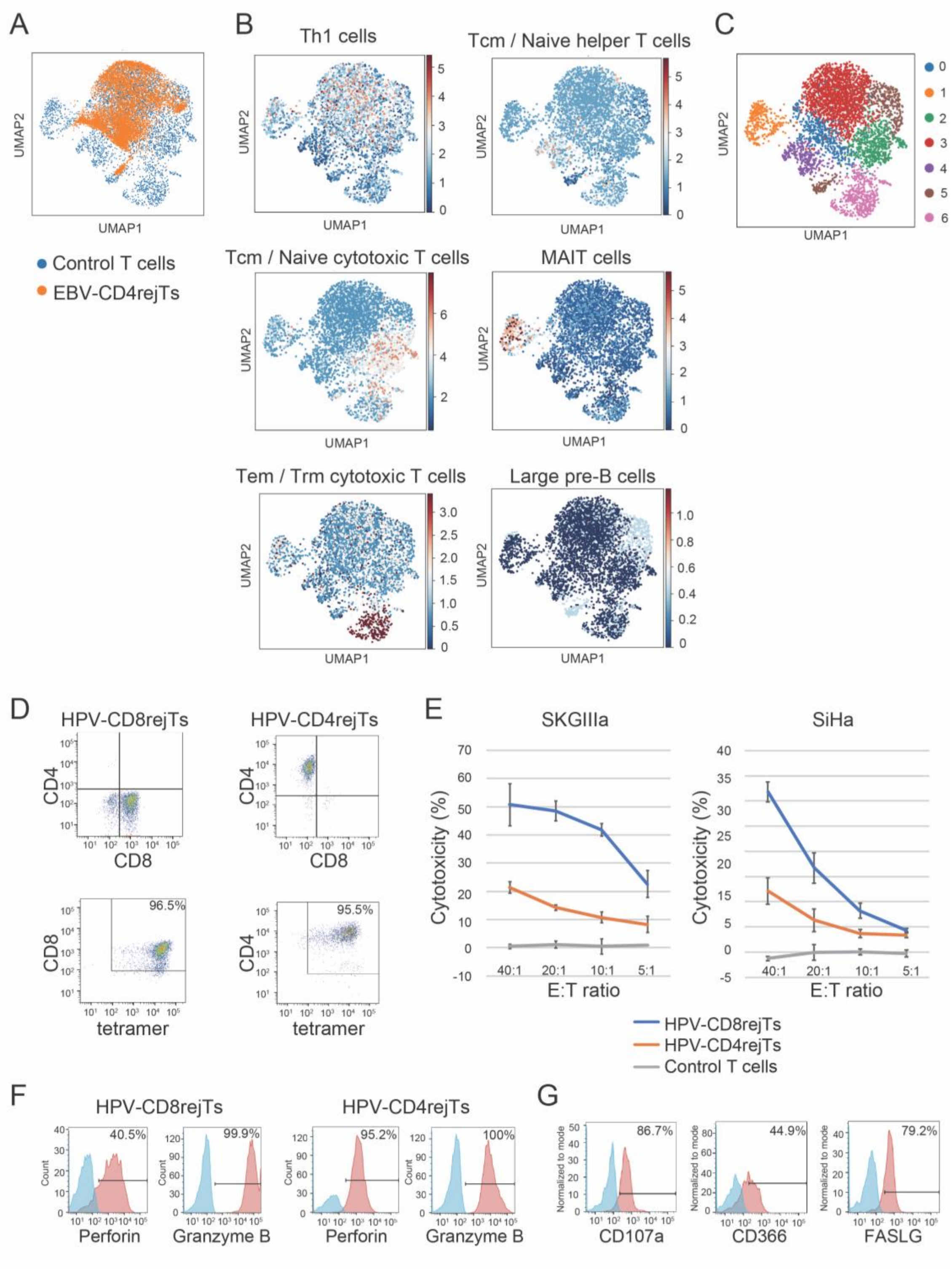


**Supplementary Figure 3. Comparison of EBV-CD4rejTs with conventional T cells**

(A) UMAP analysis of conventional T cells (control T cells) shown in blue and EBV-

CD4rejTs shown in orange. (B) Annotation of scRNA-seq data of control T cells and EBV-CD4rejTs using CellTypist automated cell type annotation for scRNA-seq datasets. (C) Annotation of scRNA-seq data of control T cells and EBV-CD4rejTs. Cluster 0: Th1 cells No. 1, cluster 1: MAIT cells, cluster 2: Tcm/Naive cytotoxic T cells, cluster 3: Th1 cells No. 2, cluster 4: Tcm/Naive helper T cells Tcm, cluster 5: large pre-B cells, cluster 6: Tem/Trm cytotoxic T cells. Tcm, central memory T cells; Tem, effector memory T cells; Trm, tissue memory T cells.

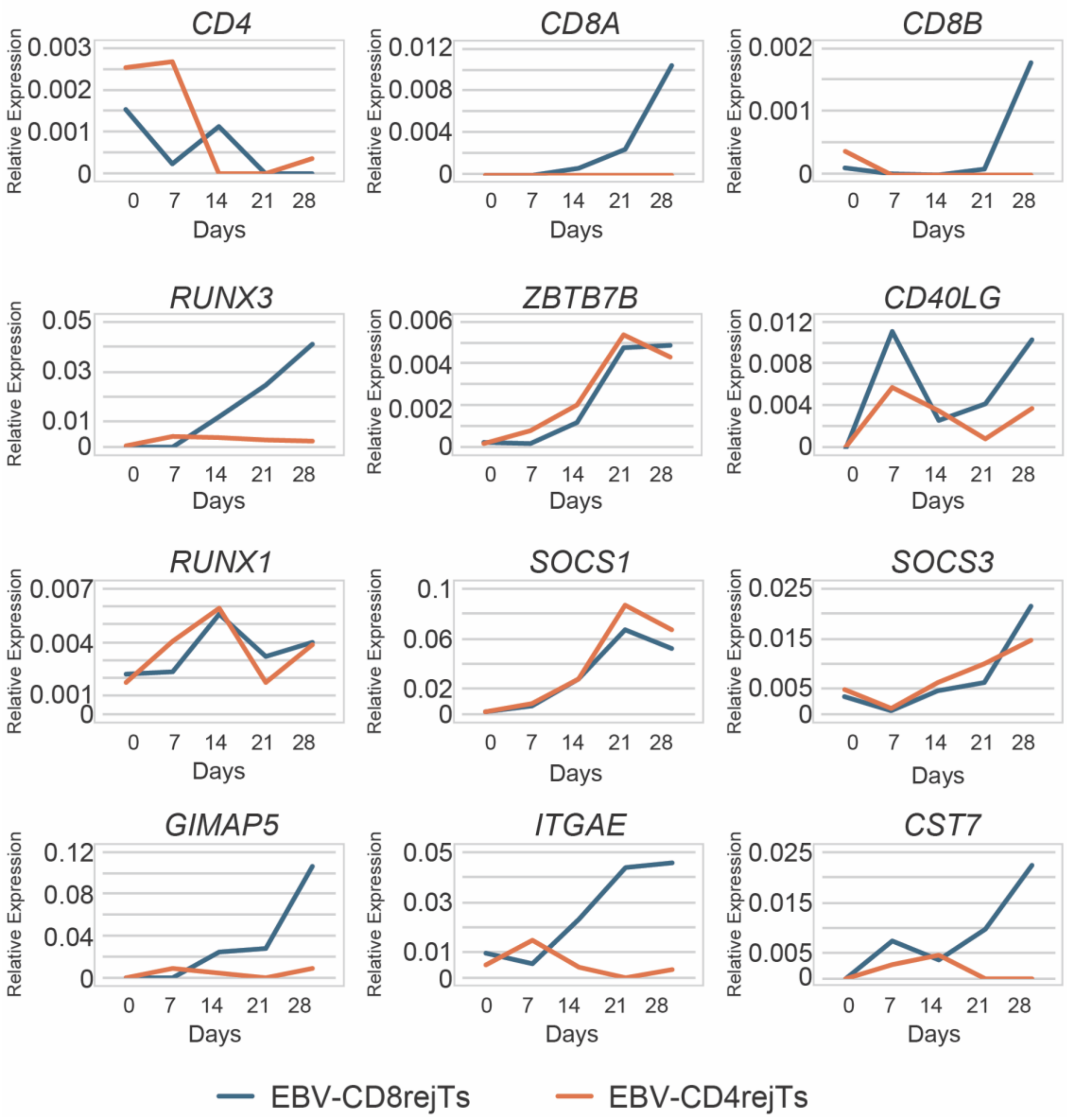


**Supplementary Figure 4. Weekly RNA-seq data of EBV-CD8rejTs and EBV-CD4rejTs during T cell differentiation**

Weekly RNA-seq analysis of genes associated with CD4/CD8 lineage choice in EBV-CD8rejTs and EBV-CD4rejTs during T cell differentiation. RNA-seq was performed after collection of iPSC-derived hematopoietic progenitor cells (Day 0) every week until day 28.

**Supplementary Table 1. Two ATL patient donors**

| Patient No. | Type | Percentage of tumor cells in peripheral blood at time of diagnosis (%) | Percentage of tumor cells in bone marrow at time of diagnosis (%) |
|---|---|---|---|
| 1 | Acute | 51 | 16.5 |
| 2 | Lymphoma | 2.4 | 1 |
